# Thermodynamic and kinetic design principles for protein aggregation inhibitors

**DOI:** 10.1101/2020.02.22.960716

**Authors:** Thomas C. T. Michaels, Andela Šarić, Georg Meisl, Gabriella T. Heller, Samo Curk, Paolo Arosio, Sara Linse, Christopher M. Dobson, Michele Vendruscolo, Tuomas P. J. Knowles

## Abstract

Understanding the mechanism of action of compounds capable of inhibiting protein aggregation is critical to the development of potential ther-apeutics against protein misfolding diseases. A fundamental challenge for progress is the range of possible target species and the disparate timescales involved, since the aggregating proteins are simultaneously the reactants, products, intermediates and catalysts of the reaction. It is a complex problem, therefore, to choose the states of the aggregating proteins that should be bound by the compounds to achieve the most potent inhibition. We present here a comprehensive kinetic theory of protein aggregation inhibition which reveals the fundamental thermodynamic and kinetic signatures characterising effective inhibitors by identifying quantitative relationships between the aggregation and binding rate constants. These results provide general physical laws to guide the design and optimisation of protein aggregation inhibitors.

---

The aggregation of peptides and proteins into amyloid fibrils is key in many phenomena ranging from the formation of functional machineries in biology to the production of novel nanomaterials. Recent interest in this process has come from the realisation that protein aggregation is intimately linked to a range of human conditions, from Alzheimer’s disease to type II diabetes.^1–6^ Inhibition of protein aggregation thus represents a major strategy for the development of effective pharmacological interventions against protein misfolding diseases.^7–24^ Traditionally, inhibition strategies against protein aggregation have focussed either on blocking the production of aggregation-prone peptides and proteins or on promoting the degradation of their amyloid products.^7^ Recent attempts have instead concentrated on altering or delaying the aggregation process itself, which is typically achieved using compounds that bind non-covalently to different types of protein species during the aggregation reaction.^8–10^ Examples of such compounds include small drug-like molecules,^23–28^ molecular chaperones,^29–31^ antibodies,^32,33^ and nanoparticles.^34^ Initially, such kinetic inhibitors were designed with the generic goal of delaying amyloid formation.^15–21^ It has now been recognised, however, that the cytotoxicity linked to amyloid formation is not attributable to a single species. Instead, the gamut of species accessible during aggregation contribute differently to toxicity.^30,35–38^ Equally importantly, it has also been established that the aggregation reaction is a complex non-linear kinetic process in which aggregating proteins can act simultaneously as the reactants, products, intermediates and catalysts of the reaction. This coupling between different aggregation steps makes it difficult to estimate the overall effects of interventions aimed at affecting the population balance of specific species.^39,43^ Thus, successful inhibition strategies must build on a detailed mechanistic understanding of the aggregation reaction network and the manner in which it is affected by inhibitors.^9^ Despite this importance, the fundamental physical principles that underlie inhibition of protein aggregation remain poorly understood. In particular, it remains challenging to establish a quantitative connection between inhibitor binding to aggregating species and the resulting inhibitory effect, and hence between thermodynamics and kinetics in protein aggregation inhibition.

To address this limitation, we present a general theory of inhibition of protein aggregation. We formulate the problem in terms of a master equation for aggregation kinetics in the presence of inhibitors that bind one or more of the aggregating species. We derive explicit integrated rate laws to such dynamic equations and show that such compounds preserve the structure of the aggregation reaction network, i.e. do not modify the final form of the analytical solution. The kinetics of inhibited aggregation can therefore be interpreted in terms of effective rate parameters, which in practice provide a clear strategy to determine the mechanistic effect of specific inhibitors from experimental data. Moreover, our framework uncovers general thermodynamic and kinetic constraints on effective inhibition by quantifying the effect of an inhibitor explicitly in terms of the aggregation rate constants and binding parameters, including binding affinities. These simple physical principles will likely facilitate the design, search and optimisation of effective inhibitors of pathological protein aggregation.

## Aggregation kinetics in the presence of an inhibitor

Dealing with the heterogeneous and transient mixture of species involved in protein aggregation has proved challenging. In the absence of inhibitors, much progress has come from applying the methodologies of chemical kinetics, well-established for simple chemical transformations, to protein aggregation processes.^39–45^ In particular, the discovery of integrated rate laws for the aggregation kinetics has provided key insights into the specific molecular steps of amyloid fibril formation from the analysis of experimental data (Fig. 1a).^39–45^ Initially, the smallest stable aggregates form directly from soluble monomers through primary nucleation.^40^ Fibrils grow by elongation, i.e. the addition of individual monomers at the aggregate ends.^40^ In many cases, including the aggregation of A*β*42,^43^ rapid proliferation of aggregates can be promoted by existing fibrils in a process termed surface-catalysed secondary nucleation.^43–48^ In this process, new fibrils nucleate through the interaction of monomers with the surfaces of existing fibrils. Fibrils thus act as catalysts of the aggregation reaction.

**Figure 1:**
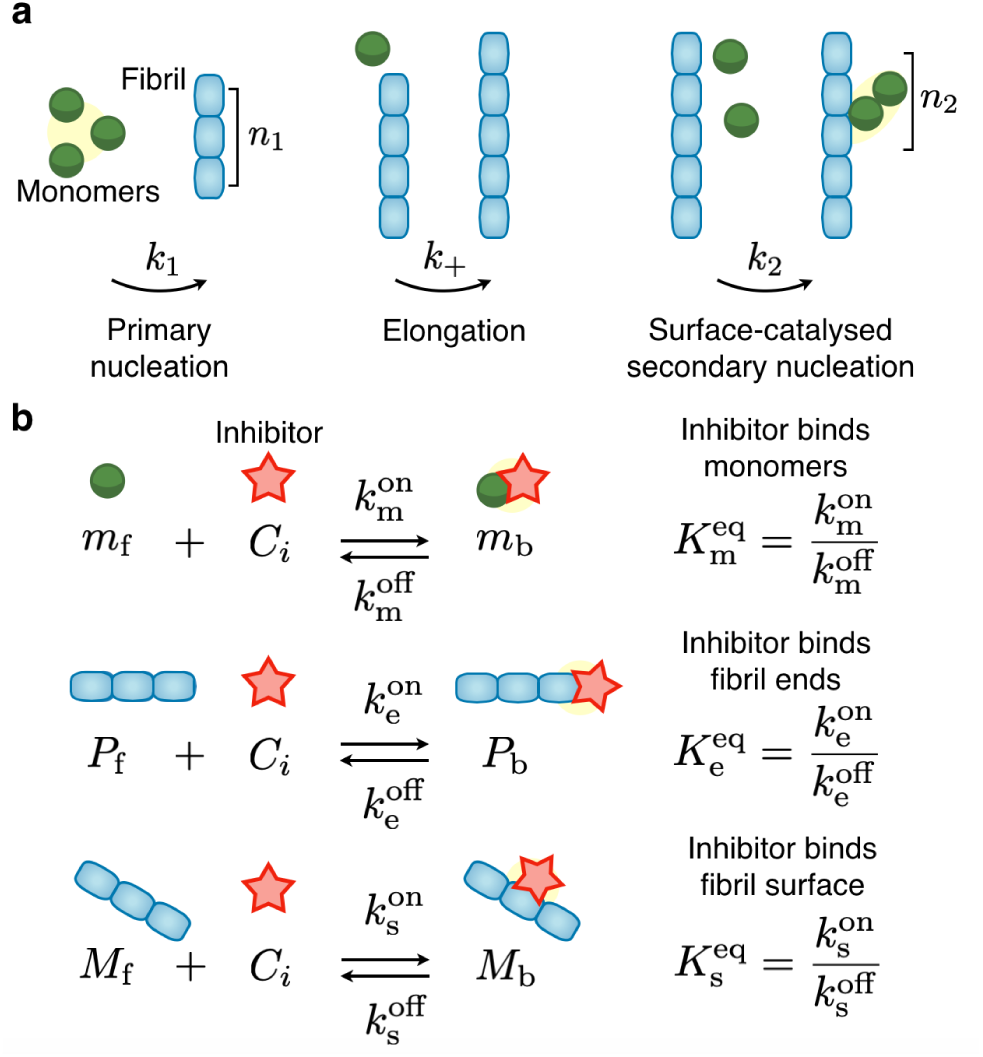
Microscopic mechanisms of protein aggregation and possible inhibition pathways. **(a)** Schematic representation of the microscopic steps of protein aggregation into fibrillar structures. **(b)** Potential target species during protein aggregation and associated binding rate constants.

Within this chemical kinetics approach, the combined effect of these different microscopic steps on the overall aggregation kinetics is captured by means of a master equation for key experimentally accessible observables, including the free monomer concentration *m*(*t*) and the number or mass concentrations of fibrils, denoted with *P*(*t*) respectively *M*(*t*) (see SI Sec. 1.1 for a derivation):^39–44^

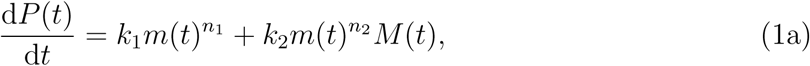

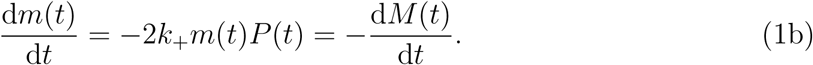

Here, *k*_+_, *k*_1_ and *k*_2_ are the rate constants for filament elongation, primary nucleation and surface-catalysed secondary nucleation. The exponents *n*_1_ and *n*_2_ are the reaction orders of primary, respectively, secondary nucleation with respect to the concentration of free monomers (SI Sec. S1.1 and Refs.^47,50–53^ for details on the physical interpretation of reaction orders). Eq. (1a) captures the total rate of formation of new aggregates by primary and secondary nucleation. Eq. (1b) describes the consumption of monomers, which occurs mainly by fibril elongation. Since monomers are either free or part of aggregates, the total concentration of monomers in the system, *m*_tot_, is conserved at all times, i.e. *m*(*t*) + *M*(*t*) = *m*_tot_.

Eqs. (1a) and (1b) yield sigmoidal-type kinetics for fibril mass concentration (Fig. 2). An explicit integrated rate law describing this behaviour has been obtained previously:^41,42^

**Figure 2:**
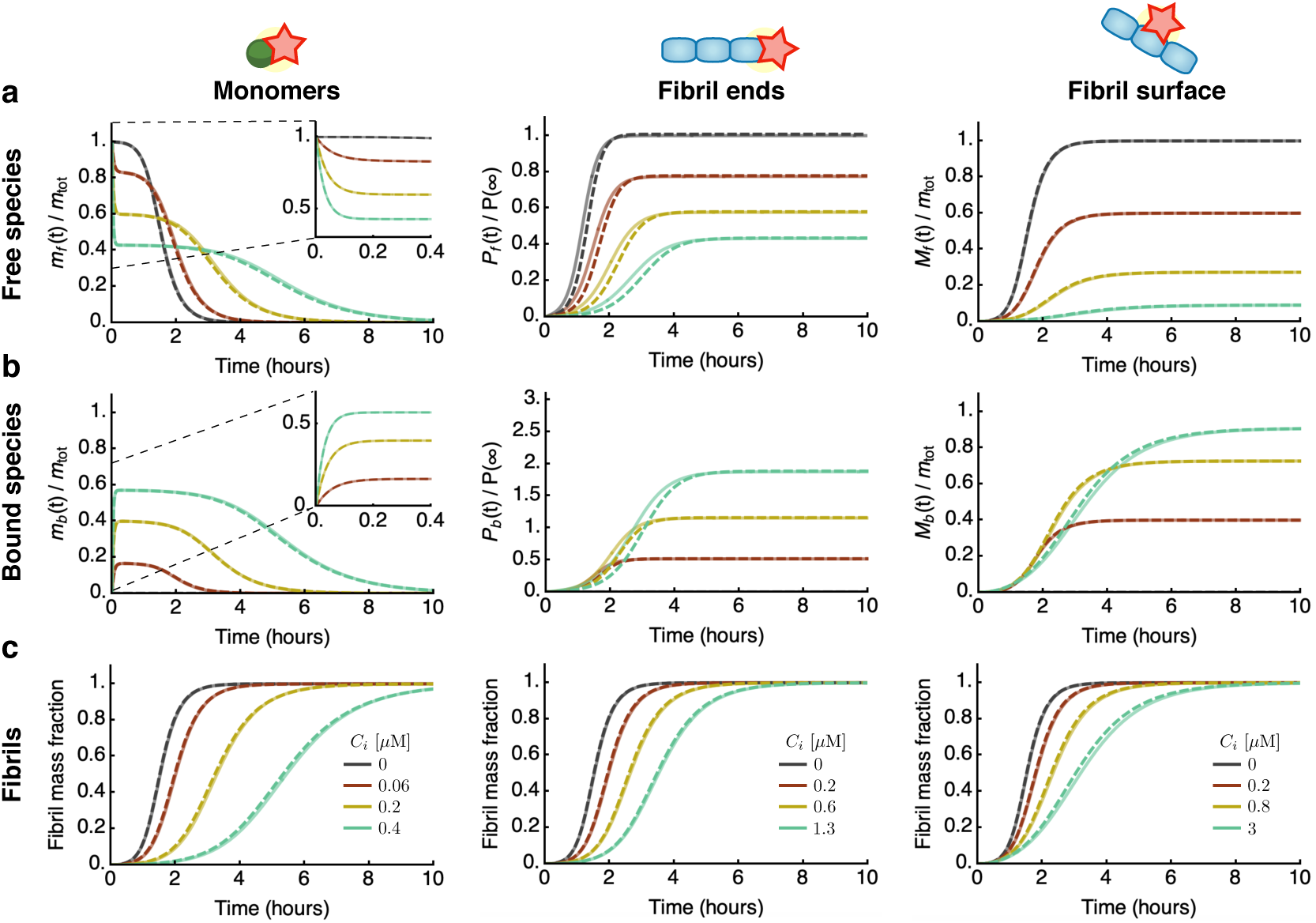
Integrated rate laws for protein aggregation in the presence of inhibitors. Characteristic kinetic profiles for **(a)** free species, **(b)** bound species, and **(c)** fibril mass concentration in the presence of an inhibitor that binds free monomers (first column), fibrils ends (middle column) or fibril surfaces (right column). Dashed lines are the analytical integrated rate laws (see SI for explicit expressions), which are in excellent agreement with numerical simulations of Eqs. (3) (solid lines). Calculation parameters: *m*_tot_ = 3*µ*M, *k*_+_ = 3 × 10^6^M^−1^s^−1^, *k*_1_ = 10^−4^M^−1^s^−1^, *k*_2_ = 8 × 10^3^M^−2^s^−1^, *n*_1_ = *n*_2_ = 2, 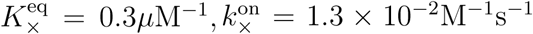, and (a) *C*_*i*_ = 0.06, 0.2, 0.4*µ*M, (b) *C*_*i*_ = 0.2, 0.6, 1.3*µ*M, (c) *C*_*i*_ = 0.2, 0.8, 3*µ*M. Black curves are without inhibitor.

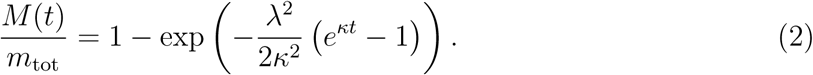

Eq. (2) reveals that only two key rate parameters control the time course of aggregation: 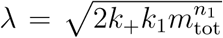 and 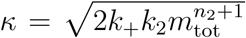. These are combined rates describing aggregate proliferation through primary and secondary nucleation, respectively. Expressions such as (2) provide a useful means to interpret protein aggregation experiments in terms of the underlying microscopic steps from global fitting of measured aggregation curves (SI Sec. S1.1 for further discussion).^54^

Within the framework of Eq. (1) an inhibitor can affect protein aggregation kinetics by binding (i) free monomers, (ii) fibril ends or (iii) fibril surface sites; these ‘species’ correspond to the three fields *m*(*t*), *P*(*t*) and *M*(*t*) (Fig. 1b and SI Sec. S1.2).^8,29^ This description may be generalised to account for inhibitor binding to additional target species, including e.g. transient oligomers.^55^ We describe the effect of inhibitors on aggregation using a master equation by introducing species for the monomer concentration, the number and mass concentrations of fibrils which are either active (“free”, subscript “f”) or deactivated due to the binding to inhibitor molecules (“bound”, subscript “b”). In our model, bound species are unable to participate in the aggregation process ^1^. We denote the binding and unbinding rates of the inhibitor to the respective species as 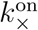, respectively, 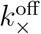, where × is a placeholder that can refer to monomers (× = m), fibrils ends (× = e), or fibril surface sites (× = s). By binding to its target, the inhibitor affects the population balance of free and bound species thereby influencing the rates of the different aggregation processes in which these species are involved.

The time course of the aggregation reaction in the presence of an inhibitor is captured by the following kinetic equations for the “free” species, as an extension of Eqs. (1):

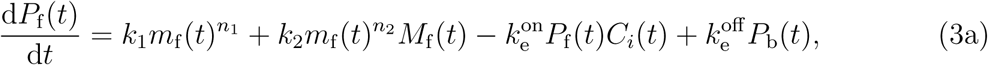

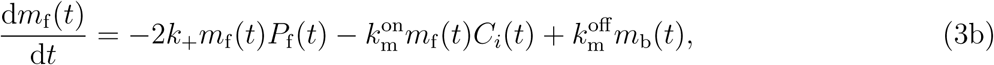

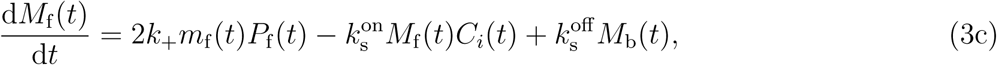

where *C*_*i*_(*t*) denotes the concentration of free inhibitor. The equations for the “bound” species read:

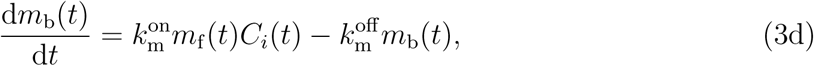

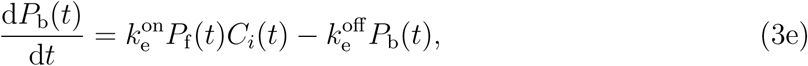

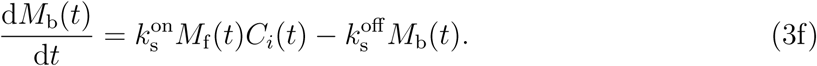

To derive Eqs. (3), we have further assumed that the amount of species (monomers, fibril ends or fibril surface sites) deactivated by the inhibitor can be calculated by considering independent binding events on the different sites. The binding rate is thus proportional to the concentration of free species and the concentration of free inhibitor *C*_*i*_(*t*).

## Integrated rate laws for inhibited aggregation kinetics

To understand how an inhibitor influences aggregation in terms of the underlying rate parameters, we derived explicit analytical solutions to Eqs. (3) by exploiting self-consistent approaches (SI Sec. S2). Strikingly, for all modes of inhibition, we find that the integrated rate law for aggregate mass concentration has the same functional form as in the absence of the inhibitor, Eq. (2) (Fig. 2, SI Eq. (S51-52)):

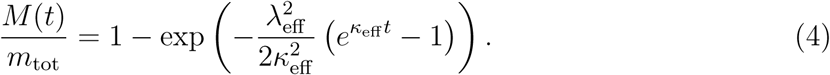

As a key result of the assumptions made here, we find that the structure of the reaction network of aggregation is preserved in the presence of inhibitors. This conclusion is true irrespective of the speed and magnitude of inhibitor binding, i.e. the rates of association and dissociation relative to the rate of aggregation. Consequently, in the presence of inhibitors the rate couplings *λ* and *κ* in Eq. (2) are replaced in Eq. (4) by effective rates, *λ*_eff_ and *κ*_eff_, which depend in characteristic ways on the kinetic parameters of aggregation and inhibitor binding. The functional dependence of *λ*_eff_ and *κ*_eff_ depends on which aggregating species are targeted (Fig. 3a,b): i) an inhibitor that binds the surface of fibrils may reduce the secondary nucleation rate, hence affecting only *κ*; ii) an inhibitor that binds fibril ends may lower the rate of elongation, affecting *λ* and *κ* equally; iii) an inhibitor that binds monomers may reduce the rates of all steps of aggregation, affecting both *λ* and *κ*, although in general by different amounts.

**Figure 3:**
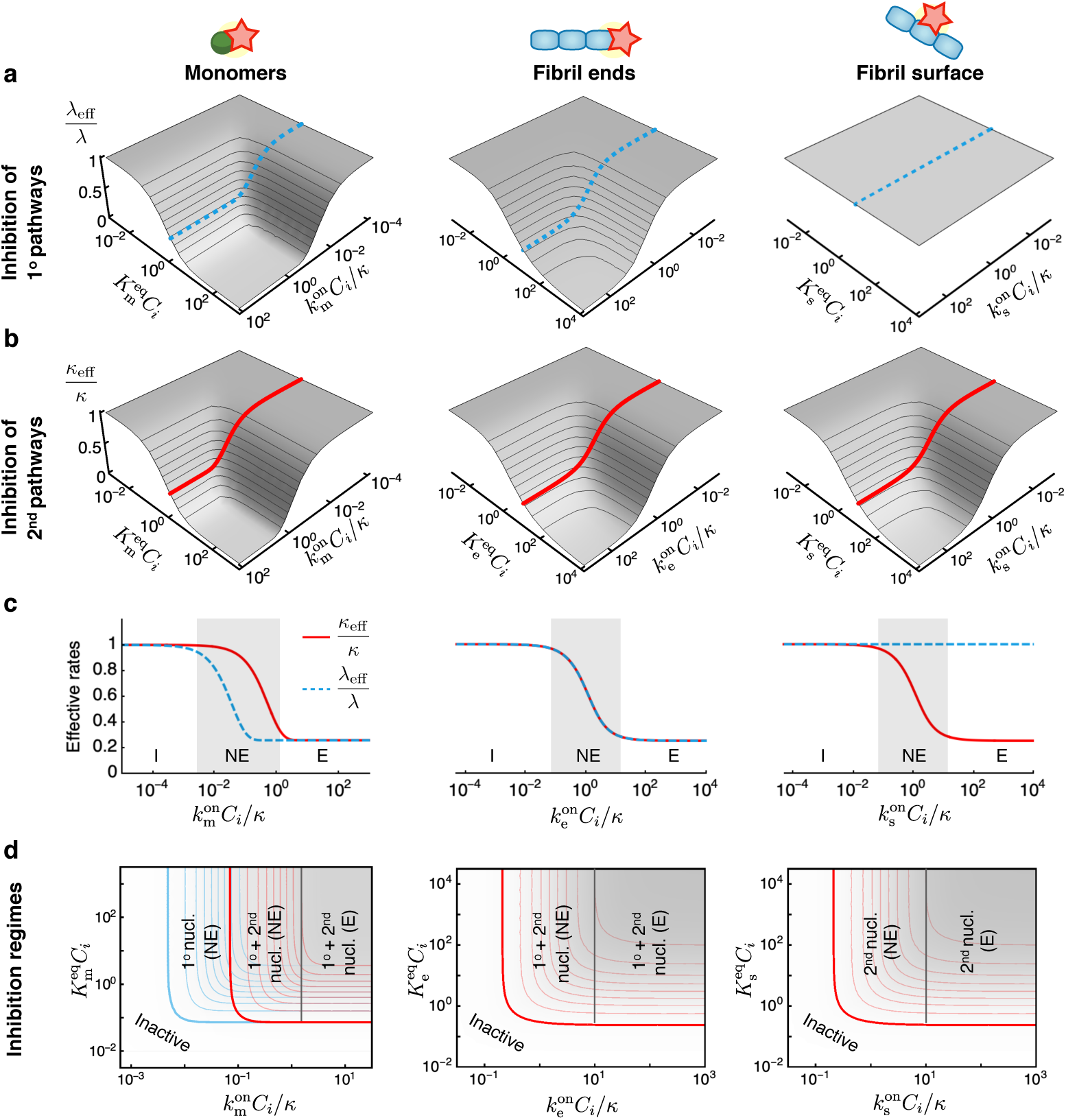
Thermodynamic and kinetic design principles for protein aggregation inhibitors. **(a-b)** Effective rates of aggregation in the presence of an inhibitor. (a) The parameter *λ*_eff_/*λ* describes the extent of inhibition on the primary aggregation pathways. (b) *κ*_eff_/*κ* describes the effect on the secondary pathways. These effective rates depend in characteristic ways on two combined parameters: a dimensionless binding rate 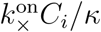 and dimensionless binding constant 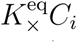. Dashed and solid lines in (a) and (b) indicate sample inhibitor-response curves as a function of 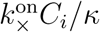 at constant 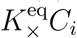. **(c)** Depending on the rate of inhibitor binding to its target, 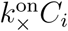, compared to the overall rate of aggregation (*κ*), we distinguish different inhibition regimes: (I) inactive 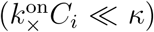, (NE) non-equilibrium inhibition 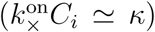, (E) equilibrium inhibition 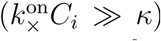. For fixed 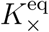, maximal inhibition is obtained in the equilibrium regime. The extent of maximal inhibition depends solely on 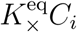. The plots in (c) correspond to the dashed and solid lines shown in (a) respectively (b). **(d)** Schematic phase diagram summarising possible inhibition regimes for an inhibitor that binds to monomers, fibril ends or fibril surfaces. These phase diagrams are top views of the plots in (a) and (b), i.e. contour plots of *λ*_eff_/*λ* (blue) and *κ*_eff_/*κ* (red) (see Fig. S3). Contour lines in (a), (b) and (c) are shown in intervals of 0.1. Plots of *κ*_eff_/*κ* and *λ*_eff_/*λ* are generated using the expressions in Table S2 for the same parameters as in Fig. 2.

## Rich inhibition phase behaviour from interplay between kinetics and thermodynamics

We have calculated explicit expressions for the effective rates *λ*_eff_ and *κ*_eff_, which are summarised in Table S2. These expressions reveal that, in the case when the inhibitor binds aggregate ends or surface sites, *λ*_eff_/*λ* and *κ*_eff_/*κ* are controlled only by two dimensionless parameters: (1) a normalised binding rate 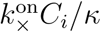 and (2) a dimensionless binding constant 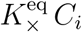, where 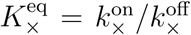 is the equilibrium binding constant. Based on these kinetic and thermodynamic parameters, we can distinguish 3 primary regimes of inhibition: (1) no inhibition 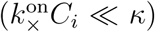, (2) non-equilibrium inhibition 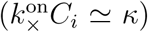, and (3) equilibrium inhibition 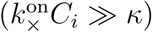. When the inhibitor binds the target species slowly compared to the characteristic timescale of aggregation (1*/κ*), the effective rates *λ*_eff_/*λ* and *κ*_eff_/*κ* are close to 1 and the inhibitor is kinetically inactive. In the opposite limit when inhibitor binding is fast compared to aggregation, we can invoke pre-equilibrium for the binding of the inhibitor to the target species (SI Secs. S1.4-5). We term this regime equilibrium inhibition, since the effective rate parameters are determined in this limit solely by the dimensionless binding constant 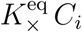 (Table S1):

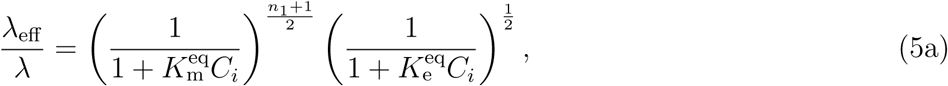

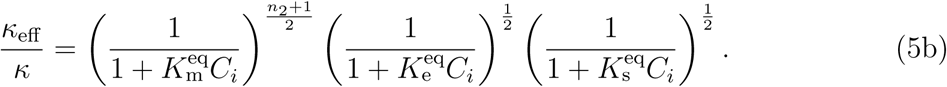

Thus, effective inhibition necessarily requires sufficiently low binding affinity 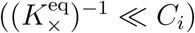 and fast binding to the target 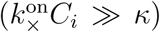. An interesting intermediate non-equilibrium inhibition regime emerges when the inhibitor binding rate to its target is comparable to the characteristic rate of aggregation *κ*. In this case, weaker inhibition is observed compared to the equilibrium limit. At fixed binding affinity, the maximum possible extent of inhibition is obtained in the equilibrium regime. Our explicit expressions for *λ*_eff_/*λ* and *κ*_eff_/*κ* interpolate smoothly between these limiting regimes (Fig. 3c).

In the case when the inhibitor binds monomers, an additional dimensionless parameter *λ/κ* emerges as the ratio of the proliferation rates through primary and secondary pathways. Since *λ* ≪ *κ*, on the basis of this parameter we can distinguish two further non-equilibrium inhibition regimes. When 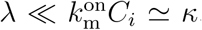, we observe a non-equilibrium inhibition regime where both primary and secondary nucleation are affected. However, when 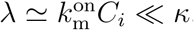, the inhibitor affects primary nucleation only, but binds monomers too slowly to be able to interfere with the secondary nucleation process (Fig. 3d, first column).

Our theory may be extended to inhibitors that bind multiple species simultaneously. In this case, we find that the effects of the individual modes of inhibition on *λ* and *κ* combine multiplicatively (see SI Sec. S2.7).

## Physical design principles for effective inhibitors and illustrative examples on experimental data

Our work carries important implications for the design of potential protein aggregation inhibitors, by revealing that specific combinations of the thermodynamic and kinetic parameters determine the effectiveness of a compound to inhibit protein aggregation. The expressions for the effective rates of inhibited aggregation, *λ*_eff_/*λ* and *κ*_eff_/*κ*, allow us to construct phase diagrams for the possible inhibition regimes (Fig. 3d and Fig. S3). These diagrams, which have kinetic and thermodynamic axes, provide precise strategies to optimise inhibition. For instance, in the case of an inhibitor that binds monomers, increasing the parameter 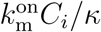 can turn an inhibitor of primary nucleation into an effective inhibitor of both primary and secondary nucleation pathways (Fig. 3d, first column). Systematically characterising different compounds on phase diagrams such as in Fig. 3d provides a novel strategy for optimising the efficacy of inhibitors of protein aggregation not only with respect to their binding affinity, but also in terms of their binding kinetics. We now illustrate this principle by considering the example of the inhibition of A*β*42 aggregation by two compounds, Brichos^30^ and 10074-G5,^56^ which selectively target different aggregating species.

Kinetic models of protein filament formation are uniquely effective for yielding information about the microscopic mechanisms of aggregation from the analysis of experimental reaction profiles.^54^ Thus, the integrated rate law obtained in this work provide a systematic framework for establishing the mechanism of action of unknown inhibitors on aggregation. We have applied this approach to experimental data on the inhibition of A*β*42 aggregation by the human Brichos domain, a molecular chaperone which has been shown to bind the surface of A*β*42 fibrils.^29,30^ The rates of binding and dissociation of Brichos to/from the surface of A*β*42 fibrils have been measured independently from the aggregation kinetics using surface plasmon resonance:^30^ 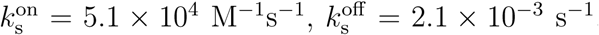, with a corresponding binding constant of 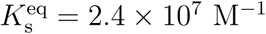. Thus, using these parameters, we can predict kinetic traces of A*β*42 aggregation in the presence of increasing concentrations of Brichos using our analytical solution without free parameters. The resulting aggregation profiles, shown in Fig. 4a, are in excellent agreement with the experimental data, highlighting the power of our approach. Interestingly, a comparison between the rate of Brichos binding and the timescale of aggregation yields 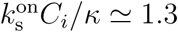 (for *C*_*i*_ = 0.3*μ*M), revealing that Brichos binds the surface of amyloid fibrils relatively slowly compared to aggregation. Indeed, the kinetic profiles predicted using the equilibrium model (Eq. (5)) do not capture the data (Fig. 4b). Moreover, a comparison between the effective *κ* and our theoretical prediction (Eq. (S76), Fig. 4c) confirms that the inhibition of A*β*42 by Brichos falls in the non-equilibrium regime (Fig. 4d).

**Figure 4:**
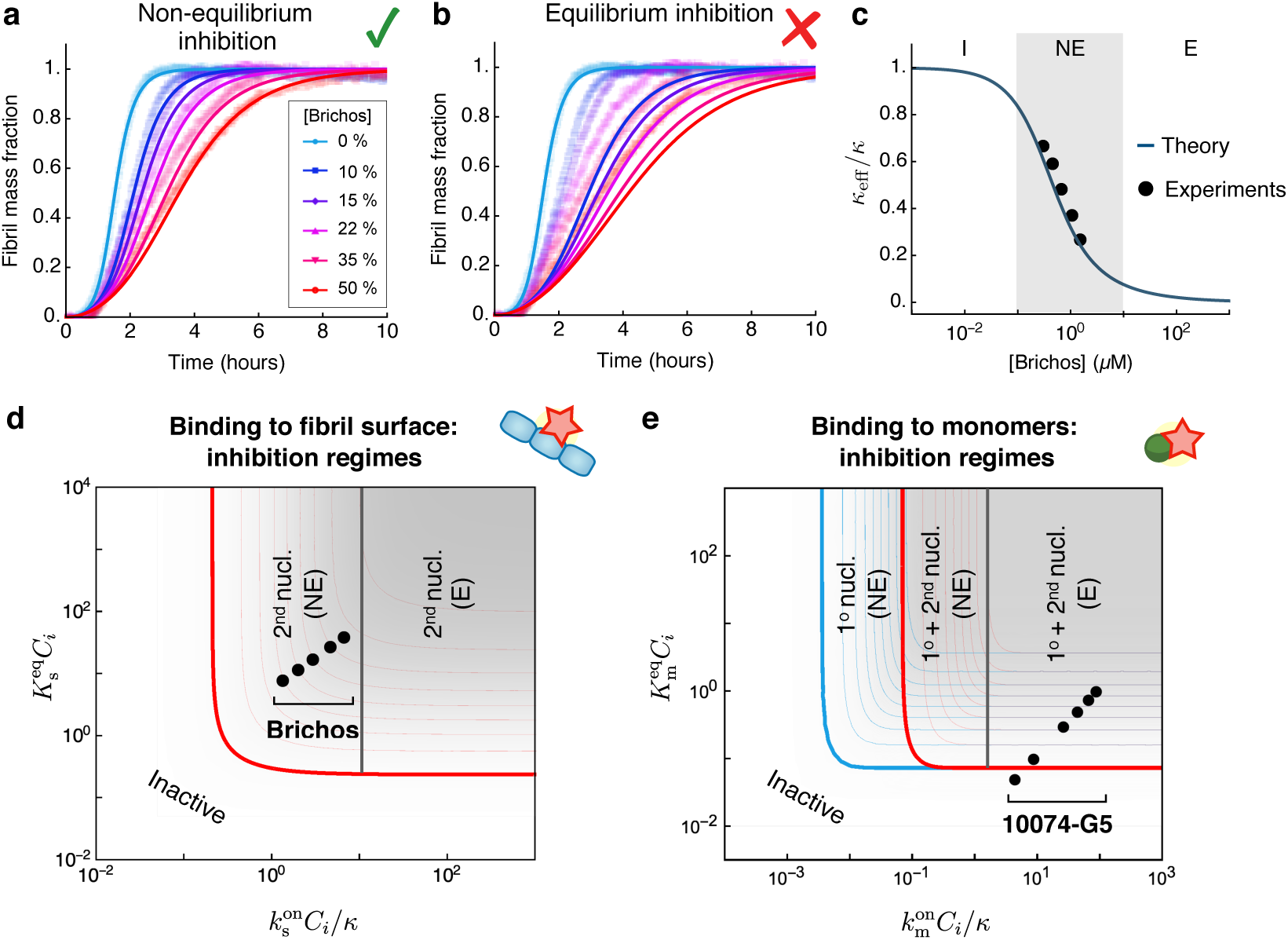
Application to the inhibition of A*β*42 aggregation. **(a)** Experimental data for the aggregation of 3*µ*M A*β*42 in the presence of increasing concentrations of Brichos (*C*_*i*_ = 0.1, 0.15, 0.22, 0.35, 0.5 A*β*42 molar equivalents). The experimental data are compared to the theoretical prediction of our integrated rate law (solid lines). The theoretical prediction for inhibited curves has no fitting parameters: the rate constants of aggregation are extracted from a fit of the aggregation curve in the absence of inhibitor (*k*_+_ = 3 × 10^6^ M^−1^s^−1^, *k*_2_ = 8.2 × 10^4^ M^−2^s−1, *k*_1_ = 1.1 × 10^−4^ M^−1^s^−1^) and the effect of inhibitor is predicted using the experimentally measured binding and dissociation rates 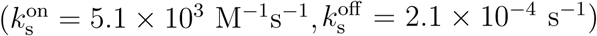.^30^ **(b)** Experimental data are compared to the theoretical prediction assuming equilibrium binding (Eq. (5)) of Brichos with binding constant 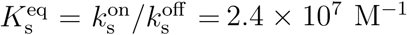. (c) Effective *κ*_eff_/*κ* as a function of Brichos concentration and comparison to our theoretical prediction (Table S2, solid line). **(d)** Location of Brichos^30^ in the phase diagram of possible inhibition regimes for an inhibitor binding fibril surface sites. The points correspond to the following concentrations of Brichos: *C*_*i*_ = 0.1, 0.15, 0.22, 0.35, 0.5 molar equivalents. **(e)** Location of 10074-G5 in the phase diagram of possible inhibition regimes for a monomer binder.^56^ The points correspond to the following inhibitor concentrations *C*_*i*_ = 1, 2, 6, 10, 15, 20*µ*M for 1*µ*M A*β*42.

As a further example we consider the inhibition of A*β*42 by the small molecule 10074-G5 (biphenyl-2-yl-(7-nitro-benzo[1,2,5]oxadiazol-4-yl)-amine), which has recently been shown to inhibit aggregation likely by binding to and sequestering monomers.^56^ The binding rate constants of 10074-G5 to monomers have been measured using bio-layer interferometry as 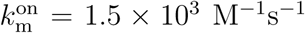 and 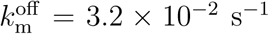,^56^ allowing us to place 10074-G5 in the phase diagram of possible inhibition regimes (Fig. 4e). This can inform future drug discovery efforts for inhibitors that bind monomers, suggesting that it is important to find compounds that bind monomers with faster on-rates.

## Summary and outlook

In this work we have developed a kinetic theory of protein aggregation inhibition. This theory offers quantitative answers to questions such as: (1) Which aggregating species should one bind to in order to suppress most effectively aggregation? (2) Which binding rates and binding affinities should be optimised? and (3) Given a lead compound, which is the best optimisation strategy? Our theory has allowed us to analyse quantitatively experimental data of aggregation inhibition outside of the conventional limiting regime where inhibitor binding is assumed to be fast compared to aggregation (equilibrium regime). Moreover, our work identifies a universal timescale (1*/κ*) to use as a ruler for developing compounds that bind specific aggregating species. These findings could guide the design and optimisation of potent inhibitors of protein aggregation by combining our theoretical framework with independent measurements of kinetics and thermodynamics of binding, which in many cases are available from experimental methods such as SPR, bio-layer interferometry and microfluidics.^29,30,56^ We also note that the principles elucidated in this study are generally applicable to linear self-assembly processes in general, and thus are not limited to inhibition of protein aggregation. For instance, these principles may be applicable to specific materials science applications, where inhibitors are used to control the physical size or other properties of aggregates.^57^

## Methods

Details of the mathematical model are available in the online version of the paper.

## Acknowledgments

We acknowledge support from Peterhouse, Cambridge (TCTM), the Swiss National Science Foundation (TCTM), the Royal Society (AS, SC), the Academy of Medical Sciences (AS), Sidney Sussex College, Cambridge (GM), Newnham College, Cambridge (GTH), the Wellcome Trust (TPJK), the Cambridge Centre for Misfolding Diseases (TPJK, MV), the BBSRC (TPJK), the Frances and Augustus Newman Foundation (TPJK). The research leading to these results has received funding from the European Research Council under the European Union’s Seventh Framework Programme (FP7/2007-2013) through the ERC grant PhysProt (agreement n° 337969).

## Competing financial interests

The authors declare no competing financial and non-financial interests.

## Data availability statement

The authors confirm that all data generated and analysed during this study are included in this published article (and its supplementary information files).

## Author contributions

All authors were involved in the design of the study; TCTM developed the theoretical model; all authors participated in interpreting the results and writing the paper.

## Footnotes

1 In reality, aggregation involves heterogeneous populations of monomers/aggregates with different aggregation propensities. Inhibitors affect the aggregation propensity of these subpopulations differently. Bound species are assumed to have a reduced propensity to participate in the aggregation process. Mathematically, this situation is equivalent to the binary formulation of Eq. (3) if binding rates are interpreted as averages over these different subpopulations.

