## Supplementary Information for "Thermodynamic and kinetic design principles for protein aggregation inhibitors"

**S3 Asymptotic solutions to aggregation kinetics with variable inhibitor concentration** **14**

### S1. Kinetic theory of protein aggregation and its inhibition

**S1.1. Kinetic equations in the absence of inhibitor.** As discussed in the main text, the time course of a protein aggregation reaction is described in terms of the underlying microscopic steps (Fig. 1a of the main text) by tracking the evolution of a few coarse-grained fields, which represent experimentally accessible observables, including

$$P(t) = \text{aggregate number concentration,} \quad \text{[S1a]}$$
$$M(t) = \text{aggregate mass concentration,} \quad \text{[S1b]}$$
$$m(t) = \text{monomer concentration.} \quad \text{[S1c]}$$

The dynamics of these three fields can be derived from a consideration of the time evolution of the entire aggregate size distribution $f(t,j)$, which describes the concentration at time $t$ of aggregates consisting of $j$ monomers. $f(t,j)$ satisfies the following master equation (1–4):

$$\frac{\partial f(t,j)}{\partial t} = 2k_+m(t)f(t,j-1) - 2k_+m(t)f(t,j) + k_1m(t)^{n_1}\delta_{j,n_1} + k_1m(t)^{n_2}\delta_{j,n_1}\sum_i if(t,j), \quad \text{[S2a]}$$

where $\delta_{i,j}$ denotes the Kronecker delta function and

$$k_1 = \text{rate constant for primary nucleation,} \quad \text{[S2b]}$$
$$k_2 = \text{rate constant for secondary nucleation,} \quad \text{[S2c]}$$
$$k_+ = \text{rate constant for aggregate elongation (growth),} \quad \text{[S2d]}$$
$$n_1 = \text{reaction order for primary nucleation,} \quad \text{[S2e]}$$
$$n_2 = \text{reaction order for secondary nucleation.} \quad \text{[S2f]}$$

A note on reaction orders for fibril nucleation: both primary and secondary nucleation of new filaments are believed to be non-classical, multi-step nucleation processes (5–7). Hence, the reaction orders $n_1$ and $n_2$ are in general not equal to the physical size of the critical nuclei, as in classical nucleation theory (5–7). Instead, the reaction orders $n_1$ and $n_2$ should be thought of as describing the dependence of the rate-limiting step of these nucleation processes on the available monomer (see e.g Ref. (7) for a discussion on the interpretation of reaction orders for protein aggregation).

The coarse-grained fields in Eq. (S1) correspond to the lowest principal moments of the aggregate size distribution $f(t, j)$

$$P(t) = \sum_j f(t, j), \quad M(t) = \sum_j j f(t, j). \tag{S3}$$

Hence, the time evolution of $P(t)$ and $M(t)$ is obtained by summation of the master equation Eq. (S2a) over $j$, yielding (1–4):

$$\frac{\mathrm{d}P(t)}{\mathrm{d}t} = k_1 m(t)^{n_1} + k_2 m(t)^{n_2} M(t), \tag{S4}$$

$$\frac{\mathrm{d}m(t)}{\mathrm{d}t} = -2k_+ m(t)P(t) - n_1 k_1 m(t)^{n_1} - n_2 k_2 m(t)^{n_2} M(t) = -\frac{\mathrm{d}M(t)}{\mathrm{d}t}. \tag{S5}$$

In general, elongation is fast compared to nucleation. Indeed, the steady-state value of the average length of aggregates can be shown to scale as $\sqrt{k_+/(k_2\, m_{\mathrm{tot}}^{n_2-1})}$ (8). Since fibrillar aggregates are typically several monomers long ($\simeq 10^4$), secondary (and hence also primary) nucleation must be slow compared to growth. This condition allows us to neglect the nucleation terms in Eq. (S5) in front of the growth term. With this simplification, we thus arrive the following set of differential equations (Eq. (1) of main text) (1–4):

$$\frac{\mathrm{d}P(t)}{\mathrm{d}t} = k_1 m(t)^{n_1} + k_2 m(t)^{n_2} M(t), \tag{S6a}$$

$$\frac{\mathrm{d}m(t)}{\mathrm{d}t} = -2k_+ m(t)P(t) = -\frac{\mathrm{d}M(t)}{\mathrm{d}t}. \tag{S6b}$$

The terms on the right-hand side of Eq. (S6a) describe the total rate of the formation of new fibrils from primary and secondary nucleation, respectively. Similarly, Eq. (S6b) describes the consumption of monomers (hence buildup of aggregate mass) through elongation. The total mass of monomers $m_{\mathrm{tot}}$ is conserved

$$m_{\mathrm{tot}} = m(t) + M(t). \tag{S7}$$

An analytical solution to the aggregation kinetics in the absence of an inhibitor has been obtained previously using self-consistent approaches and reads (1, 2, 4):

$$\frac{M(t)}{m_{\mathrm{tot}}} = 1 - \exp\left(-\frac{\lambda^2}{2\kappa^2}\left(e^{\kappa t} - 1\right)\right), \tag{S8}$$

where

$$\lambda = \sqrt{2k_+ k_1 m_{\mathrm{tot}}^{n_1}}, \tag{S9}$$

$$\kappa = \sqrt{2k_+ k_2 m_{\mathrm{tot}}^{n_2+1}} \tag{S10}$$

are effective aggregate proliferation rates of aggregates through primary and secondary nucleation, respectively. An alternative analytical solution, which is more accurate than Eq. (S8) for $n_2 \geq 1$, is given by (1, 3, 4)

$$\frac{M(t)}{m_{\mathrm{tot}}} = 1 - \left(\frac{B_+ + C_+}{B_+ + C_+ e^{\kappa t}} \frac{B_- + C_+ e^{\kappa t}}{B_- + C_+}\right)^{\frac{k_\infty^2}{\kappa \tilde{k}_\infty}} e^{-k_\infty t}, \tag{S11a}$$

where

$$C_\pm = \pm\frac{\lambda^2}{2\kappa^2}, \tag{S11b}$$

$$k_\infty = \kappa\sqrt{\frac{2}{n_2(n_2+1)} + \frac{2\lambda^2}{n_1\kappa^2}}, \tag{S11c}$$

$$\tilde{k}_\infty = \sqrt{k_\infty^2 - 4C_+C_-\kappa^2}, \tag{S11d}$$

$$B_\pm = \frac{k_\infty \pm \tilde{k}_\infty}{2\kappa}. \tag{S11e}$$

Also in this case, the solution is dependent on $\lambda$ and $\kappa$.

**S1.2. Modes of inhibition.** To understand how the presence of inhibitor molecules affects the aggregation dynamics described by Eq. (S6a) and Eq. (S6b), we consider in detail the possible modes of inhibition when a drug-like small molecule is incorporated into the dynamics of aggregation. In general, we distinguish three main scenarios for how the inhibitor molecule can interfere with the aggregation process (Fig. 1b of main text):

1. *Binding to monomers* – The first possibility for the inhibitor to influence the aggregation process is by reversibly binding to the monomers. Through reversible binding and unbinding, the monomers can be activated or deactivated. Deactivated monomers have a reduced propensity to participate to the aggregation process. We denote the rate constants for binding to and unbinding from the monomers as $k_{\mathrm{m}}^{\mathrm{on}}$ and $k_{\mathrm{m}}^{\mathrm{off}}$, respectively. The ratio $k_{\mathrm{m}}^{\mathrm{on}}/k_{\mathrm{m}}^{\mathrm{off}} = K_{\mathrm{m}}^{\mathrm{eq}}$ is the equilibrium constant for monomer binding.

2. *Binding to fibril ends* – Another possibility is that inhibitor molecules block the ends of fibrils, thereby preventing them from growing by recruiting free monomers from solution. The rate constants for binding to and unbinding from the fibril ends are indicated respectively as $k_{\mathrm{e}}^{\mathrm{on}}$ and $k_{\mathrm{e}}^{\mathrm{off}}$, and the associated equilibrium binding constant is $k_{\mathrm{e}}^{\mathrm{on}}/k_{\mathrm{e}}^{\mathrm{off}} = K_{\mathrm{e}}^{\mathrm{eq}}$. For simplicity, we assume that inhibitor binding/unbinding occur with the same rates at both fibril ends; our approach could in principle be generalised to account for different binding and dissociation rates at and from both fibril ends.

3. *Binding to fibril surface* – A third relevant possibility to consider is the binding of inhibitor molecules to the catalytic surface of existing fibrils. Inhibitor molecules bind the surface with rate constant $k_{\mathrm{s}}^{\mathrm{on}}$, thereby potentially blocking the autocatalytic cycle of surface-catalyzed secondary nucleation. Inhibitor molecules can unbind from the fibril surface with rate constant $k_{\mathrm{s}}^{\mathrm{off}}$, thereby allowing aggregates to catalyze again the formation of new aggregates on their surface. The equilibrium constant for binding to the surface of fibrils is $k_{\mathrm{s}}^{\mathrm{on}}/k_{\mathrm{s}}^{\mathrm{off}} = K_{\mathrm{s}}^{\mathrm{eq}}$.

**S1.3. Kinetic equations in the presence of inhibitor.** Depending on the specific chemical characteristics of the inhibitor molecule under consideration, all or only a subgroup of the various modes of inhibition described in the previous section could be active. In the most general scenario, protein aggregation kinetics in the presence of an inhibitor is captured by the following set of coupled differential equations, as an extension of Eq. (S6a) and Eq. (S6b) (Eq. (3) of main text):

$$\frac{\mathrm{d}P_{\mathrm{f}}(t)}{\mathrm{d}t} = k_1 m_{\mathrm{f}}(t)^{n_1} + k_2 m_{\mathrm{f}}(t)^{n_2} M_{\mathrm{f}}(t) - k_{\mathrm{e}}^{\mathrm{on}} P_{\mathrm{f}}(t) C_i(t) + k_{\mathrm{e}}^{\mathrm{off}} P_{\mathrm{b}}(t), \tag{S12a}$$

$$\frac{\mathrm{d}M_{\mathrm{f}}(t)}{\mathrm{d}t} = 2k_+ m_{\mathrm{f}}(t) P_{\mathrm{f}}(t) - k_{\mathrm{s}}^{\mathrm{on}} M_{\mathrm{f}}(t) C_i(t) + k_{\mathrm{s}}^{\mathrm{off}} M_{\mathrm{b}}(t), \tag{S12b}$$

$$\frac{\mathrm{d}m_{\mathrm{f}}(t)}{\mathrm{d}t} = -2k_+ m_{\mathrm{f}}(t) P_{\mathrm{f}}(t) - k_{\mathrm{m}}^{\mathrm{on}} m_{\mathrm{f}}(t) C_i(t) + k_{\mathrm{m}}^{\mathrm{off}} m_{\mathrm{b}}(t), \tag{S12c}$$

$$\frac{\mathrm{d}P_{\mathrm{b}}(t)}{\mathrm{d}t} = k_{\mathrm{e}}^{\mathrm{on}} P_{\mathrm{f}}(t) C_i(t) - k_{\mathrm{e}}^{\mathrm{off}} P_{\mathrm{b}}(t), \tag{S12d}$$

$$\frac{\mathrm{d}M_{\mathrm{b}}(t)}{\mathrm{d}t} = k_{\mathrm{s}}^{\mathrm{on}} M_{\mathrm{f}}(t) C_i(t) - k_{\mathrm{s}}^{\mathrm{off}} M_{\mathrm{b}}(t), \tag{S12e}$$

$$\frac{\mathrm{d}m_{\mathrm{b}}(t)}{\mathrm{d}t} = k_{\mathrm{m}}^{\mathrm{on}} m_{\mathrm{f}}(t) C_i(t) - k_{\mathrm{m}}^{\mathrm{off}} m_{\mathrm{b}}(t), \tag{S12f}$$

$$\frac{\mathrm{d}C_i(t)}{\mathrm{d}t} = -\frac{\mathrm{d}P_{\mathrm{b}}(t)}{\mathrm{d}t} - \frac{\mathrm{d}M_{\mathrm{b}}(t)}{\mathrm{d}t} - \frac{\mathrm{d}m_{\mathrm{b}}(t)}{\mathrm{d}t}, \tag{S12g}$$

where

$$P_{\mathrm{f}}(t) = \text{free aggregate number concentration,} \tag{S12h}$$
$$P_{\mathrm{b}}(t) = \text{bound aggregate number concentration,} \tag{S12i}$$
$$M_{\mathrm{f}}(t) = \text{free aggregate mass concentration,} \tag{S12j}$$
$$M_{\mathrm{b}}(t) = \text{bound aggregate mass concentration,} \tag{S12k}$$
$$m_{\mathrm{f}}(t) = \text{free monomer concentration,} \tag{S12l}$$
$$m_{\mathrm{b}}(t) = \text{bound monomer concentration,} \tag{S12m}$$
$$C_i(t) = \text{(free) inhibitor concentration,} \tag{S12n}$$
$$k_{\mathrm{e}}^{\mathrm{on}} = \text{on rate constant for inhibitor binding to aggregate ends,} \tag{S12o}$$
$$k_{\mathrm{e}}^{\mathrm{off}} = \text{off rate constant for inhibitor binding to aggregate ends,} \tag{S12p}$$
$$k_{\mathrm{s}}^{\mathrm{on}} = \text{on rate constant for inhibitor binding to aggregate surface,} \tag{S12q}$$
$$k_{\mathrm{s}}^{\mathrm{off}} = \text{off rate constant for inhibitor binding to aggregate surface,} \tag{S12r}$$
$$k_{\mathrm{m}}^{\mathrm{on}} = \text{on rate constant for inhibitor binding to monomers,} \tag{S12s}$$
$$k_{\mathrm{m}}^{\mathrm{off}} = \text{off rate constant for inhibitor binding to monomers.} \tag{S12t}$$

Eq. (S12a)-Eq. (S12e) must be coupled to the conservation of total protein mass $m_{\mathrm{tot}}$, which implies:

$$m_{\mathrm{tot}} = m_{\mathrm{f}}(t) + m_{\mathrm{b}}(t) + M_{\mathrm{f}}(t) + M_{\mathrm{b}}(t). \tag{S13}$$

***Total monomer and aggregate concentrations – effective kinetic equations.*** It is useful to introduce total aggregate number, aggregate mass and monomer concentrations as

$$P(t) = P_\mathrm{f}(t) + P_\mathrm{b}(t), \quad \text{[S14a]}$$
$$M(t) = M_\mathrm{f}(t) + M_\mathrm{b}(t), \quad \text{[S14b]}$$
$$m(t) = m_\mathrm{f}(t) + m_\mathrm{b}(t), \quad \text{[S14c]}$$

which satisfy the following equations

$$\frac{\mathrm{d}P(t)}{\mathrm{d}t} = k_1 m_\mathrm{f}(t)^{n_1} + k_2 m_\mathrm{f}(t)^{n_2} M_\mathrm{f}(t), \quad \text{[S14d]}$$
$$\frac{\mathrm{d}m(t)}{\mathrm{d}t} = -2k_+ m_\mathrm{f}(t) P_\mathrm{f}(t) = -\frac{\mathrm{d}M(t)}{\mathrm{d}t}. \quad \text{[S14e]}$$

Eq. (S14d) and Eq. (S14e) highlight the origin of inhibition: compared to uninhibited kinetics, Eq. (S6a) and Eq. (S6b), free (instead of total) monomer, aggregate number and aggregate mass concentrations appear on the right hand side of the kinetic equations Eq. (S14d) and Eq. (S14e). Thus, different microscopic events of aggregation (elongation, primary and secondary nucleation) are inhibited depending to which of the "aggregate species" $P$, $M$ or $m$ the inhibitor binds. As we will see in Sec. S2, the speed of inhibitor binding to the targeted species also determines the efficacy of its inhibitory action.

**S1.4. Fast inhibitor binding (equilibrium binding regime).** Important simplifications emerge in the limit of fast binding of the inhibitor to monomers and aggregates (the meaning of "fast" can be quantified rigorously using asymptotic analysis, see Sec. S2 and Eq. (S62)). We term this regime equilibrium inhibition regime. In this case, time variations of bound species can be approximately set equal to zero in Eq. (S12d)-Eq. (S12f)

$$\frac{\mathrm{d}P_\mathrm{b}(t)}{\mathrm{d}t} \simeq \frac{\mathrm{d}M_\mathrm{b}(t)}{\mathrm{d}t} \simeq \frac{\mathrm{d}m_\mathrm{b}(t)}{\mathrm{d}t} \simeq 0. \quad \text{[S15]}$$

This implies

$$\frac{\mathrm{d}C_i(t)}{\mathrm{d}t} \simeq 0, \quad \text{[S16]}$$

i.e. the inhibitor concentration is approximately constant. This procedure yields simple relationships that link the concentrations of free and bound material as

$$m_\mathrm{f}(t) = K_\mathrm{m}^\mathrm{eq} m_\mathrm{b}(t), \quad \text{[S17a]}$$
$$P_\mathrm{f}(t) = K_\mathrm{e}^\mathrm{eq} P_\mathrm{b}(t), \quad \text{[S17b]}$$
$$M_\mathrm{f}(t) = K_\mathrm{s}^\mathrm{eq} M_\mathrm{b}(t), \quad \text{[S17c]}$$

where

$$K_\mathrm{m}^\mathrm{eq} = \frac{k_\mathrm{m}^\mathrm{on}}{k_\mathrm{m}^\mathrm{off}}, \quad \text{[S17d]}$$
$$K_\mathrm{e}^\mathrm{eq} = \frac{k_\mathrm{e}^\mathrm{on}}{k_\mathrm{e}^\mathrm{off}}, \quad \text{[S17e]}$$
$$K_\mathrm{s}^\mathrm{eq} = \frac{k_\mathrm{s}^\mathrm{on}}{k_\mathrm{s}^\mathrm{off}} \quad \text{[S17f]}$$

are the equilibrium constants for inhibitor binding to monomers, fibril ends or fibril surface, respectively. Using Eq. (S14) we obtain relationships between the amount of free and total material, as:

$$m_\mathrm{f}(t) = \frac{m(t)}{1 + K_\mathrm{m}^\mathrm{eq} C_i^\mathrm{tot}}, \quad \text{[S18]}$$
$$P_\mathrm{f}(t) = \frac{P(t)}{1 + K_\mathrm{e}^\mathrm{eq} C_i^\mathrm{tot}}, \quad \text{[S19]}$$
$$M_\mathrm{f}(t) = \frac{M(t)}{1 + K_\mathrm{s}^\mathrm{eq} C_i^\mathrm{tot}}. \quad \text{[S20]}$$

Thus, Eq. (S14d) and Eq. (S14e) become:

$$\frac{\mathrm{d}M(t)}{\mathrm{d}t} = 2k_+ \left(\frac{m(t)}{1 + K_\mathrm{m}^\mathrm{eq} C_i^\mathrm{tot}}\right)\left(\frac{P(t)}{1 + K_\mathrm{e}^\mathrm{eq} C_i^\mathrm{tot}}\right), \quad \text{[S21]}$$
$$\frac{\mathrm{d}P(t)}{\mathrm{d}t} = k_1 \left(\frac{m(t)}{1 + K_\mathrm{m}^\mathrm{eq} C_i^\mathrm{tot}}\right)^{n_1} + k_2 \left(\frac{m(t)}{1 + K_\mathrm{m}^\mathrm{eq} C_i^\mathrm{tot}}\right)^{n_2}\left(\frac{M(t)}{1 + K_\mathrm{s}^\mathrm{eq} C_i^\mathrm{tot}}\right). \quad \text{[S22]}$$

**Table S1.** Effective couplings (rate constants) of the various steps of aggregation in the equilibrium regime.

| Targeted species | Monomers | Aggregate ends | Fibril surface |
|---|---|---|---|
| Inhibited steps | | | |
| Primary nucleation | ✓ | ✗ | ✗ |
| Secondary nucleation | ✓ | ✗ | ✓ |
| Elongation | ✓ | ✓ | ✗ |
| Effective rate constants (equilibrium inhibition) | $\frac{k_1^{\text{eff}}}{k_1} = \left(\frac{1}{1+K_{\text{m}}^{\text{eq}}C_i}\right)^{n_1}$<br>$\frac{k_2^{\text{eff}}}{k_2} = \left(\frac{1}{1+K_{\text{m}}^{\text{eq}}C_i}\right)^{n_2}$<br>$\frac{k_+^{\text{eff}}}{k_+} = \frac{1}{1+K_{\text{m}}^{\text{eq}}C_i}$ | $\frac{k_1^{\text{eff}}}{k_1} = 1$<br>$\frac{k_2^{\text{eff}}}{k_2} = 1$<br>$\frac{k_+^{\text{eff}}}{k_+} = \frac{1}{1+K_{\text{e}}^{\text{eq}}C_i}$ | $\frac{k_1^{\text{eff}}}{k_1} = 1$<br>$\frac{k_2^{\text{eff}}}{k_2} = \frac{1}{1+K_{\text{s}}^{\text{eq}}C_i}$<br>$\frac{k_+^{\text{eff}}}{k_+} = 1$ |

Eq. (S21) and Eq. (S22) are equivalent to the kinetic equations in the absence of inhibitor, Eq. (S6a) and Eq. (S6b), but the kinetic parameters are replaced by "effective" rate constants that depend on the inhibitor concentration (Table S1).

$$\frac{k_+^{\text{eff}}}{k_+} = \left(\frac{1}{1+K_{\text{m}}^{\text{eq}}C_i^{\text{tot}}}\right)\left(\frac{1}{1+K_{\text{e}}^{\text{eq}}C_i^{\text{tot}}}\right), \qquad \text{[S23]}$$

$$\frac{k_1^{\text{eff}}}{k_1} = \left(\frac{1}{1+K_{\text{m}}^{\text{eq}}C_i^{\text{tot}}}\right)^{n_1}, \qquad \text{[S24]}$$

$$\frac{k_2^{\text{eff}}}{k_2} = \left(\frac{1}{1+K_{\text{m}}^{\text{eq}}C_i^{\text{tot}}}\right)^{n_2}\left(\frac{1}{1+K_{\text{s}}^{\text{eq}}C_i^{\text{tot}}}\right). \qquad \text{[S25]}$$

The time course of aggregate mass concentration in the presence of an inhibitor in the fast binding limit can therefore be obtained by replacing the rate parameters in Eq. (S8) by Eq. (S23)-Eq. (S25), i.e.

$$\frac{M(t)}{m_{\text{tot}}} = 1 - \exp\left(-\frac{\lambda_{\text{eff}}^2}{2\kappa_{\text{eff}}^2}\left(e^{\kappa_{\text{eff}}t}-1\right)\right), \qquad \text{[S26]}$$

where the effective kinetic parameters are given by:

$$\frac{\lambda_{\text{eff}}}{\lambda} = \left(\frac{1}{1+K_{\text{m}}^{\text{eq}}C_i^{\text{tot}}}\right)^{\frac{n_1+1}{2}}\left(\frac{1}{1+K_{\text{e}}^{\text{eq}}C_i^{\text{tot}}}\right)^{\frac{1}{2}}, \qquad \text{[S27]}$$

$$\frac{\kappa_{\text{eff}}}{\kappa} = \left(\frac{1}{1+K_{\text{m}}^{\text{eq}}C_i^{\text{tot}}}\right)^{\frac{n_2+1}{2}}\left(\frac{1}{1+K_{\text{e}}^{\text{eq}}C_i^{\text{tot}}}\right)^{\frac{1}{2}}\left(\frac{1}{1+K_{\text{s}}^{\text{eq}}C_i^{\text{tot}}}\right)^{\frac{1}{2}}. \qquad \text{[S28]}$$

When $n_2 \geq 1$, Eq. (S11a) may be used to describe inhibited aggregation kinetics, where $\lambda$ and $\kappa$ are replaced by $\lambda_{\text{eff}}$ and $\kappa_{\text{eff}}$, respectively.

**S1.5. Scaling argument to determine inhibition regimes.** In Sec. S1.4, we have assumed "fast" inhibitor binding. We can determine the relevant timescale that differentiates the different inhibition regimes using a simple scaling argument. We illustrate this idea for an inhibitor that binds aggregate ends. A rigorous timescale analysis based on matched asymptotics is given in Sec. S2. Successful inhibition requires binding to be sufficiently fast; to quantify the meaning of "fast" in this case, we need to compare the terms $k_{\text{e}}^{\text{on}}P_{\text{f}}(t)C_i(t)$ and $k_2 m_{\text{f}}(t)^{n_2}M_{\text{f}}(t)$ in Eq. (S12). Equating these two terms, using the fact that aggregate number concentration scales as $P \simeq \kappa/(2k_+)$ (8), yields

$$k_2 m_{\text{tot}}^{n_2+1} \simeq \frac{\kappa}{2k_+}k_{\text{e}}^{\text{on}}C_i \quad \Rightarrow \quad k_{\text{e}}^{\text{on}}C_i \simeq \kappa. \qquad \text{[S29]}$$

This simple argument shows that the relevant timescale for comparing inhibitor binding is $1/\kappa$. In general, $1/\kappa$ emerges as the key timescale to which the inhibitor binding rate $k_\times^{\text{on}}C_i$ to the target $\times = \text{m,e,s}$ must be compared.

### S2. Asymptotic solutions to aggregation kinetics in the presence of an inhibitor

We now discuss the mathematical details associated with the derivation of analytical solutions to the aggregation kinetics in the presence of an inhibitor, Eq. (4) of the main text.

**S2.1. Separation of timescales.** We construct an analytical solution to Eq. (S12) using singular perturbation approaches rooted in asymptotic analysis (9). Asymptotic analysis is useful in the context of protein aggregation kinetics, since there is a separation of timescales between primary nucleation (which is very slow) and the subsequent growth of aggregates (which is comparatively very fast). This timescale separation is formalised in terms of the the following parameter

$$\varepsilon = \frac{k_1 m_{\mathrm{tot}}^{n_1-2}}{2k_+} \ll 1, \tag{S30}$$

which is typically much less than unity. For example, typical values for $\varepsilon$ in the case of the amyloid-$\beta$ peptide are $\varepsilon = 5 \times 10^{-11}$ for A$\beta$42 (3) or $\varepsilon = 3 \times 10^{-12}$ for A$\beta$40 (10). Physically, a small $\varepsilon$ is necessary to ensure that the aggregates that are formed during the reaction are long. If nucleation is slow compared to growth, few nuclei will form which can grow very long. On the contrary, when nucleation is fast compared to growth, many nuclei can form; fewer monomers, however, will be available for growth, causing aggregates to be shorter on average (see discussion after Eq. (S5)).

**S2.2. Perturbation expansion.** Since $\varepsilon$ can be considered as small perturbation parameter, we construct a perturbation solution. To this end, it is convenient to rewrite Eq. (S12) in dimensionless form first:

$$\frac{\mathrm{d}\bar{P}_\mathrm{f}(\tau)}{\mathrm{d}\tau} = \varepsilon\, \bar{m}_\mathrm{f}(\tau)^{n_1} + \nu_2\, \bar{m}_\mathrm{f}(\tau)^{n_2}\Big(1 - \bar{m}_\mathrm{f}(\tau) - \bar{m}_\mathrm{b}(\tau) - \bar{M}_\mathrm{b}(\tau)\Big) - \beta_\mathrm{e}\, \bar{P}_\mathrm{f}(\tau) + \alpha_\mathrm{e}\, \bar{P}_\mathrm{b}(\tau), \tag{S31a}$$

$$\frac{\mathrm{d}\bar{m}_\mathrm{f}(\tau)}{\mathrm{d}\tau} = -\bar{m}_\mathrm{f}(\tau)\bar{P}_\mathrm{f}(\tau) - \beta_\mathrm{m}\, \bar{m}_\mathrm{f}(\tau) + \alpha_\mathrm{m}\, \bar{m}_\mathrm{b}(\tau), \tag{S31b}$$

$$\frac{\mathrm{d}\bar{m}_\mathrm{b}(\tau)}{\mathrm{d}\tau} = \beta_\mathrm{m}\, \bar{m}_\mathrm{f}(\tau) - \alpha_\mathrm{m}\, \bar{m}_\mathrm{b}(\tau), \tag{S31c}$$

$$\frac{\mathrm{d}\bar{P}_\mathrm{f}(\tau)}{\mathrm{d}\tau} = \beta_\mathrm{e}\, \bar{P}_\mathrm{f}(\tau) - \alpha_\mathrm{e}\, \bar{P}_\mathrm{b}(\tau), \tag{S31d}$$

$$\frac{\mathrm{d}\bar{M}_\mathrm{b}(\tau)}{\mathrm{d}\tau} = \beta_\mathrm{s}\Big(1 - \bar{m}_\mathrm{f}(\tau) - \bar{m}_\mathrm{b}(\tau) - \bar{M}_\mathrm{b}(\tau)\Big) - \alpha_\mathrm{s}\, \bar{M}_\mathrm{b}(\tau), \tag{S31e}$$

where we have eliminated the equation for $M_\mathrm{f}(t)$ using conservation of mass, Eq. (S13), we have defined

$$\bar{P}_\mathrm{f} = \frac{P_\mathrm{f}(t)}{m_\mathrm{tot}}, \tag{S31f}$$

$$\bar{P}_\mathrm{b} = \frac{P_\mathrm{b}(t)}{m_\mathrm{tot}}, \tag{S31g}$$

$$\bar{M}_\mathrm{f} = \frac{M_\mathrm{f}(t)}{m_\mathrm{tot}}, \tag{S31h}$$

$$\bar{M}_\mathrm{b} = \frac{M_\mathrm{b}(t)}{m_\mathrm{tot}}, \tag{S31i}$$

$$\bar{m}_\mathrm{f} = \frac{m_\mathrm{f}(t)}{m_\mathrm{tot}}, \tag{S31j}$$

$$\bar{m}_\mathrm{b} = \frac{m_\mathrm{b}(t)}{m_\mathrm{tot}}, \tag{S31k}$$

$$\tau = 2k_+ m_\mathrm{tot} t, \tag{S31l}$$

and we have introduced the following dimensionless parameters:

$$\nu_2 = \frac{k_2 m_{\mathrm{tot}}^{n_2-1}}{2k_+}, \tag{S31m}$$

$$\beta_\mathrm{e} = \frac{k_\mathrm{e}^{\mathrm{on}} C_i^{\mathrm{tot}}}{2k_+ m_\mathrm{tot}}, \tag{S31n}$$

$$\beta_\mathrm{s} = \frac{k_\mathrm{s}^{\mathrm{on}} C_i^{\mathrm{tot}}}{2k_+ m_\mathrm{tot}}, \tag{S31o}$$

$$\beta_\mathrm{m} = \frac{k_\mathrm{m}^{\mathrm{on}} C_i^{\mathrm{tot}}}{2k_+ m_\mathrm{tot}}, \tag{S31p}$$

$$\alpha_\mathrm{e} = \frac{k_\mathrm{e}^{\mathrm{off}}}{2k_+ m_\mathrm{tot}}, \tag{S31q}$$

$$\alpha_\mathrm{s} = \frac{k_\mathrm{s}^{\mathrm{off}}}{2k_+ m_\mathrm{tot}}, \tag{S31r}$$

$$\alpha_\mathrm{m} = \frac{k_\mathrm{m}^{\mathrm{off}}}{2k_+ m_\mathrm{tot}}. \tag{S31s}$$

A perturbation series solution of Eq. (S31) can now be constructed as

$$\bar{P}_\mathrm{f}(\tau) = \bar{P}_\mathrm{f}^{(0)}(\tau) + \varepsilon\,\bar{P}_\mathrm{f}^{(1)}(\tau) + \mathcal{O}(\varepsilon^2), \quad \text{[S32a]}$$

$$\bar{P}_\mathrm{b}(\tau) = \bar{P}_\mathrm{b}^{(0)}(\tau) + \varepsilon\,\bar{P}_\mathrm{b}^{(1)}(\tau) + \mathcal{O}(\varepsilon^2), \quad \text{[S32b]}$$

$$\bar{M}_\mathrm{f}(\tau) = \bar{M}_\mathrm{f}^{(0)}(\tau) + \varepsilon\,\bar{M}_\mathrm{f}^{(1)}(\tau) + \mathcal{O}(\varepsilon^2), \quad \text{[S32c]}$$

$$\bar{M}_\mathrm{b}(\tau) = \bar{M}_\mathrm{b}^{(0)}(\tau) + \varepsilon\,\bar{M}_\mathrm{b}^{(1)}(\tau) + \mathcal{O}(\varepsilon^2), \quad \text{[S32d]}$$

$$\bar{m}_\mathrm{f}(\tau) = \bar{m}_\mathrm{f}^{(0)}(\tau) + \varepsilon\,\bar{m}_\mathrm{f}^{(1)}(\tau) + \mathcal{O}(\varepsilon^2), \quad \text{[S32e]}$$

$$\bar{m}_\mathrm{b}(\tau) = \bar{m}_\mathrm{b}^{(0)}(\tau) + \varepsilon\,\bar{m}_\mathrm{b}^{(1)}(\tau) + \mathcal{O}(\varepsilon^2). \quad \text{[S32f]}$$

***S2.2.1. $\mathcal{O}(\varepsilon^0)$ solution – initial layer dynamics.*** After inserting the perturbation expansions Eq. (S32) in Eq. (S12) and then collecting terms for each order of $\varepsilon$, we arrive at the equations at order $\varepsilon^0$:

$$\frac{\mathrm{d}\bar{P}_\mathrm{f}^{(0)}(\tau)}{\mathrm{d}\tau} = \nu_2\,\bar{m}_\mathrm{f}^{(0)}(\tau)^{n_2}\left(1 - \bar{m}_\mathrm{f}^{(0)}(\tau) - \bar{m}_\mathrm{b}^{(0)}(\tau) - \bar{M}_\mathrm{b}^{(0)}(\tau)\right) - \beta_\mathrm{e}\,\bar{P}_\mathrm{f}^{(0)}(\tau) + \alpha_\mathrm{e}\,\bar{P}_\mathrm{b}^{(0)}(\tau), \quad \text{[S33a]}$$

$$\frac{\mathrm{d}\bar{m}_\mathrm{f}^{(0)}(\tau)}{\mathrm{d}\tau} = -\bar{m}_\mathrm{f}^{(0)}(\tau)\bar{P}_\mathrm{f}^{(0)}(\tau) - \beta_\mathrm{m}\,\bar{m}_\mathrm{f}^{(0)}(\tau) + \alpha_\mathrm{m}\,\bar{m}_\mathrm{b}^{(0)}(\tau), \quad \text{[S33b]}$$

$$\frac{\mathrm{d}\bar{m}_\mathrm{b}^{(0)}(\tau)}{\mathrm{d}\tau} = \beta_\mathrm{m}\,\bar{m}_\mathrm{f}^{(0)}(\tau) - \alpha_\mathrm{m}\,\bar{m}_\mathrm{b}^{(0)}(\tau), \quad \text{[S33c]}$$

$$\frac{\mathrm{d}\bar{P}_\mathrm{b}^{(0)}(\tau)}{\mathrm{d}\tau} = \beta_\mathrm{e}\,\bar{P}_\mathrm{f}^{(0)}(\tau) - \alpha_\mathrm{e}\,\bar{P}_\mathrm{b}^{(0)}(\tau), \quad \text{[S33d]}$$

$$\frac{\mathrm{d}\bar{M}_\mathrm{b}^{(0)}(\tau)}{\mathrm{d}\tau} = \beta_\mathrm{s}\left(1 - \bar{m}_\mathrm{f}^{(0)}(\tau) - \bar{m}_\mathrm{b}^{(0)}(\tau) - \bar{M}_\mathrm{b}^{(0)}(\tau)\right) - \alpha_\mathrm{s}\,\bar{M}_\mathrm{b}^{(0)}(\tau). \quad \text{[S33e]}$$

Applying the initial conditions $\bar{m}_\mathrm{f}^{(0)}(0) = 1$ and $\bar{P}_\mathrm{f}^{(0)}(0) = \bar{P}_\mathrm{b}^{(0)}(0) = \bar{M}_\mathrm{f}^{(0)}(0) = \bar{M}_\mathrm{b}^{(0)}(0) = \bar{m}_\mathrm{b}^{(0)}(0) = 0$, we see immediately that the solution to the $O(\varepsilon^0)$ equation, Eq. (S33), is:

$$\bar{P}_\mathrm{f}^{(0)} \equiv 0, \quad \text{[S34a]}$$

$$\bar{P}_\mathrm{b}^{(0)} \equiv 0, \quad \text{[S34b]}$$

$$\bar{M}_\mathrm{f}^{(0)} \equiv 0, \quad \text{[S34c]}$$

$$\bar{M}_\mathrm{b}^{(0)} \equiv 0, \quad \text{[S34d]}$$

$$\bar{m}_\mathrm{f}^{(0)} = 1 - \frac{\beta_\mathrm{m}}{\alpha_\mathrm{m} + \beta_\mathrm{m}}\left(1 - e^{-(\alpha_\mathrm{m}+\beta_\mathrm{m})\tau}\right), \quad \text{[S34e]}$$

$$\bar{m}_\mathrm{b}^{(0)} = \frac{\beta_\mathrm{m}}{\alpha_\mathrm{m} + \beta_\mathrm{m}}\left(1 - e^{-(\alpha_\mathrm{m}+\beta_\mathrm{m})\tau}\right). \quad \text{[S34f]}$$

Hence, due to to the separation of timescales between nucleation and growth, the dynamics of the system evolves initially through a rapid phase of equilibration, where the inhibitor binds monomers but no aggregates form at leading order in $\varepsilon$. Note that during this initial phase, the total monomer concentration $\bar{m}^{(0)} = \bar{m}_\mathrm{f}^{(0)} + \bar{m}_\mathrm{b}^{(0)} = 1$ is constant at leading order in $\varepsilon$. During the initial layer phase, which is the temporal equivalent of a boundary layer, the initial value of the monomer concentration relaxes quickly to the equilibrium value $\frac{\beta_\mathrm{m}}{\alpha_\mathrm{m}+\beta_\mathrm{m}}$ before any aggregation occurs (see Figs. S1 and Fig. 2 of the main text).

***S2.2.2. $\mathcal{O}(\varepsilon^1)$ solution – slow manifold.*** After this initial, rapid phase of monomer redistribution through inhibitor binding, the system enters a slower phase of dynamics where, at leading order in $\varepsilon$, the system stays on the slow manifold

$$\bar{m}_\mathrm{f}(\tau) = \frac{\alpha_\mathrm{m}}{\alpha_\mathrm{m} + \beta_\mathrm{m}}\bar{m}(\tau) \quad \text{[S35]}$$

at all times. This relationship, valid in the slow manifold, is verified against numerical integration of Eq. (S12) in Fig. S1d.

To obtain a solution valid for the slow manifold, we collect terms of order $\varepsilon^1$ in our perturbation expansion of Eq. (S31).

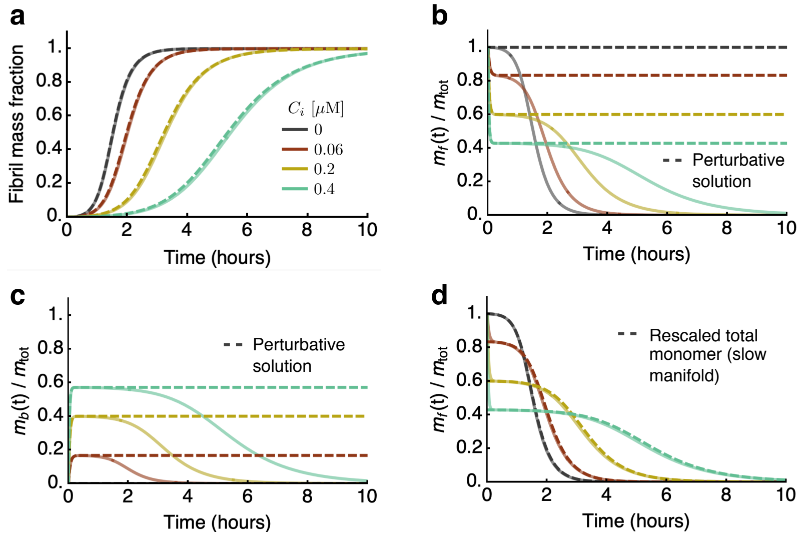

**Fig. S1.** (a) Time course of aggregate mass concentration in the presence of increasing concentrations of an inhibitor of that binds monomers calculated using numerical integration of the master equation Eq. (S12) (solid lines) and our analytical solution (dashed line). (b-c) Free and bound monomer concentrations. In the initial layer phase, there is rapid binding of the inhibitor to the free monomers (dashed lines indicate the $\mathcal{O}(\varepsilon^0)$ perturbation solution Eq. (S34)); no aggregation occurs during this phase. A slow manifold phase follows, where the both the free and bound monomer concentrations decrease slowly due to aggregation. (d) In the slow manifold, there is a relationship linking the free monomer concentration to the total monomer concentration Eq. (S35). This relationship is verified here numerically through integration of the master equation Eq. (S12) (solid line: free monomer concentration, dashed line: Eq. (S35)). Calculation parameters are the same as in the left column of Fig. 2 of the main text.

Using Eq. (S34), we arrive at the following first order equations:

$$\frac{\mathrm{d}\bar{P}_\mathrm{f}^{(1)}(\tau)}{\mathrm{d}\tau} = \bar{m}_\mathrm{f}^{(0)}(\tau)^{n_1} + \nu_2\, \bar{m}_\mathrm{f}^{(0)}(\tau)^{n_2}\Big( - \bar{m}_\mathrm{f}^{(1)}(\tau) - \bar{m}_\mathrm{b}^{(1)}(\tau) - \bar{M}_\mathrm{b}^{(1)}(\tau)\Big) - \beta_\mathrm{e}\, \bar{P}_\mathrm{f}^{(1)}(\tau) + \alpha_\mathrm{e}\, \bar{P}_\mathrm{b}^{(1)}(\tau) \quad \text{[S36a]}$$

$$\frac{\mathrm{d}\bar{m}_\mathrm{f}^{(1)}(\tau)}{\mathrm{d}\tau} = -\bar{m}_\mathrm{f}^{(0)}(\tau)\bar{P}_\mathrm{f}^{(1)}(\tau) - \beta_\mathrm{m}\, \bar{m}_\mathrm{f}^{(1)}(\tau) + \alpha_\mathrm{m}\, \bar{m}_\mathrm{b}^{(1)}(\tau), \quad \text{[S36b]}$$

$$\frac{\mathrm{d}\bar{m}_\mathrm{b}^{(1)}(\tau)}{\mathrm{d}\tau} = \beta_\mathrm{m}\, \bar{m}_\mathrm{f}^{(1)}(\tau) - \alpha_\mathrm{m}\, \bar{m}_\mathrm{b}^{(1)}(\tau), \quad \text{[S36c]}$$

$$\frac{\mathrm{d}\bar{P}_\mathrm{b}^{(1)}(\tau)}{\mathrm{d}\tau} = \beta_\mathrm{e}\, \bar{P}_\mathrm{f}^{(1)}(\tau) - \alpha_\mathrm{e}\, \bar{P}_\mathrm{b}^{(1)}(\tau), \quad \text{[S36d]}$$

$$\frac{\mathrm{d}\bar{M}_\mathrm{b}^{(1)}(\tau)}{\mathrm{d}\tau} = \beta_\mathrm{s}\Big( - \bar{m}_\mathrm{f}^{(1)}(\tau) - \bar{m}_\mathrm{b}^{(1)}(\tau) - \bar{M}_\mathrm{b}^{(1)}(\tau)\Big) - \alpha_\mathrm{s}\, \bar{M}_\mathrm{b}^{(1)}(\tau), \quad \text{[S36e]}$$

where, to match with the $O(\varepsilon^0)$ solution, we set

$$\bar{m}_\mathrm{f}^{(0)} = \frac{\alpha_\mathrm{m}}{\alpha_\mathrm{m} + \beta_\mathrm{m}} =: \mu_0. \quad \text{[S37]}$$

Eq. (S37) can also be written as

$$\mu_0 = \frac{1}{1 + \beta_\mathrm{m}/\alpha_\mathrm{m}} = \frac{1}{1 + K_\mathrm{m}^\mathrm{eq} C_i^\mathrm{tot}}. \quad \text{[S38]}$$

Eq. (S36) is therefore a set of coupled linear differential equations, which we can write in matrix form as

$$\frac{\mathrm{d}\boldsymbol{x}}{\mathrm{d}t} = \boldsymbol{A}\boldsymbol{x} + \boldsymbol{b} \quad \text{[S39]}$$

or, explicitly,

$$\frac{\mathrm{d}}{\mathrm{d}\tau}\begin{pmatrix} \bar{P}_\mathrm{f}^{(1)} \\ \bar{m}_\mathrm{f}^{(1)} \\ \bar{m}_\mathrm{b}^{(1)} \\ \bar{P}_\mathrm{b}^{(1)} \\ \bar{M}_\mathrm{b}^{(1)} \end{pmatrix} = \underbrace{\begin{pmatrix} -\beta_\mathrm{e} & -\nu_2\mu_0^{n_2} & -\nu_2\mu_0^{n_2} & \alpha_\mathrm{e} & -\nu_2\mu_0^{n_2} \\ -\mu_0 & -\beta_\mathrm{m} & \alpha_\mathrm{m} & 0 & 0 \\ 0 & \beta_\mathrm{m} & -\alpha_\mathrm{m} & 0 & 0 \\ \beta_\mathrm{e} & 0 & 0 & -\alpha_\mathrm{e} & 0 \\ 0 & -\beta_\mathrm{s} & -\beta_\mathrm{s} & 0 & -(\alpha_\mathrm{s} + \beta_\mathrm{s}) \end{pmatrix}}_{=\boldsymbol{A}} \underbrace{\begin{pmatrix} \bar{P}_\mathrm{f}^{(1)} \\ \bar{m}_\mathrm{f}^{(1)} \\ \bar{m}_\mathrm{b}^{(1)} \\ \bar{P}_\mathrm{b}^{(1)} \\ \bar{M}_\mathrm{b}^{(1)} \end{pmatrix}}_{=\boldsymbol{x}} + \underbrace{\begin{pmatrix} \mu_0^{n_1} \\ 0 \\ 0 \\ 0 \\ 0 \end{pmatrix}}_{=\boldsymbol{b}}. \quad \text{[S40]}$$

The solution to Eq. (S40) with initial condition $\boldsymbol{x}(0) = \boldsymbol{0}$ can be written as

$$\boldsymbol{x}(\tau) = \int_0^\tau e^{\boldsymbol{A}(\tau-\tau')}\boldsymbol{b}\, \mathrm{d}\tau'. \quad \text{[S41]}$$

To determine the exponential matrix of $\boldsymbol{A}$, we diagonalise $\boldsymbol{A}$. Let $x_1, x_2, \cdots$ denote the eigenvalues of $\boldsymbol{A}$. For simplicity, we order these eigenvalues in such a way that $x_1$ is the largest, real, positive eigenvalue (the order of the other eigenvalues does not matter for our calculation). That such an eigenvalue exists will be shown below. If we now construct a matrix $\boldsymbol{U}$ in such a way that the first column of $\boldsymbol{U}$ is the eigenvector to the eigenvalue $x_1$, the second column of $\boldsymbol{U}$ is the eigenvector to the eigenvalue $x_2$, and so on, then we can write the matrix $\boldsymbol{A}$ as follows:

$$\boldsymbol{A} = \boldsymbol{U}\boldsymbol{D}\boldsymbol{U}^{-1}, \tag{S42}$$

where

$$\boldsymbol{D} = \begin{pmatrix} x_1 & & \\ & x_2 & \\ & & \ddots \end{pmatrix} \tag{S43}$$

is a diagonal matrix consisting of the different eigenvalues of $\boldsymbol{A}$. Hence, the solution to Eq. (S40) can be written as

$$\boldsymbol{x}(\tau) = \int_0^\tau e^{\boldsymbol{A}(\tau-\tau')}\boldsymbol{b}\, d\tau' = \boldsymbol{U}\left(\int_0^\tau e^{\boldsymbol{D}(\tau-\tau')}d\tau'\right)\boldsymbol{U}^{-1}\boldsymbol{b} = \boldsymbol{U}\begin{pmatrix} \frac{(e^{x_1\tau}-1)}{x_1} & & \\ & \frac{(e^{x_2\tau}-1)}{x_2} & \\ & & \ddots \end{pmatrix}\boldsymbol{U}^{-1}\boldsymbol{b}. \tag{S44}$$

The solution is a sum of exponentials; the longer-time behavior is going to be dominated by the fastest growing exponential term, which corresponds to the largest positive eigenvalue $x_1$. Hence, after a rapid phase of adjustment, aggregate concentrations increase exponentially with time with the characteristic, dominant multiplication rate $x_1$.

The accuracy of the exponential solution Eq. (S44) against numerical integration of the master equation Eq. (S12) is shown in Fig. S2a for the example of an inhibitor binding fibril surface sites. While this solution captures the dynamics of the system in the early stages, it does not capture saturation at late times. We now use a self-consistent approach (1, 2, 11) to construct, from this exponentially growing solution, an accurate expression for the aggregate mass which captures the sigmoidal nature of kinetics (see Ref. (11) for an introduction to self-consistent methods to protein aggregation kinetics).

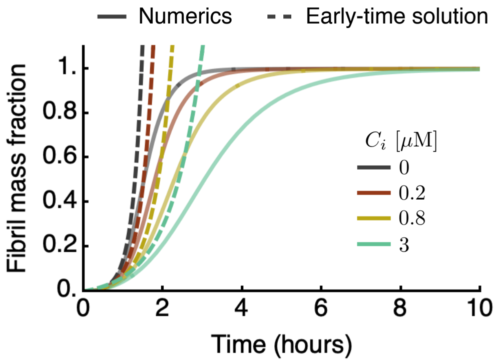

**Fig. S2.** Time course of aggregate mass concentration in the presence of increasing concentrations of an inhibitor of secondary nucleation calculated using the early-time exponential perturbation solution Eq. (S44) (dashed lines) is compared to numerical integration of the master equation Eq. (S12) (solid lines). Calculation parameters are the same as in the right column of Fig. 2 of the main text.

**S2.3. Self-consistent solution.** To construct a self-consistent solution for the total aggregate mass concentration, we start from Eq. (S14e), which using Eq. (S35) can be re-written as

$$\frac{\mathrm{d}m(t)}{\mathrm{d}t} = -2k_+ m_\mathrm{f}(t)P_\mathrm{f}(t) = -2k_+\mu_0\, m(t)P_\mathrm{f}(t) = -\frac{\mathrm{d}M(t)}{\mathrm{d}t}. \tag{S45}$$

The formal solution to Eq. (S45) is thus

$$\frac{M(t)}{m_\mathrm{tot}} = 1 - \exp\left(-2k_+\mu_0 \int_0^t P_\mathrm{f}(s)ds\right). \tag{S46}$$

Finding an explicit expression for the aggregate mass thus requires knowledge of the time evolution of the free aggregate number concentration $P_\mathrm{f}(t)$. At leading order, we can substitute the perturbation series expansion solution for $P_\mathrm{f}$, yielding:

$$\begin{aligned}\bar{M}(\tau) &= 1 - \exp\left(-\mu_0 \int_0^\tau \bar{P}_\mathrm{f}(\tau')d\tau'\right) \simeq 1 - \exp\left(-\mu_0 \int_0^\tau \left(\bar{P}_\mathrm{f}^{(0)}(\tau') + \varepsilon\, \bar{P}_\mathrm{f}^{(1)}(\tau') + \cdots\right)d\tau'\right) \\ &= 1 - \exp\left(-\mu_0\, \varepsilon \int_0^\tau \bar{P}_\mathrm{f}^{(1)}(\tau')d\tau'\right).\end{aligned} \tag{S47}$$

Hence, we need an expression for $\bar{P_{\rm f}}^{(1)}$. This can be obtained from Eq. (S44) as:

$$\bar{P_{\rm f}}^{(1)}(\tau) = x_1(\tau) = \mu_0^{n_1} \sum_{j=1}^{n} U_{1j} \frac{(e^{x_j \tau} - 1)}{x_j} \left(U^{-1}\right)_{j1}, \tag{S48}$$

where $U_{ij}$ denotes the $ij$-th component of the matrix $\boldsymbol{U}$. Recall that $x_1$ is the largest positive, real eigenvalue of $\boldsymbol{A}$. Keeping only the dominant exponential growing term $e^{x_1\tau}$ (in front of slower growing or decaying terms $e^{x_2\tau}, \cdots$), we find

$$\bar{P_{\rm f}}^{(1)}(\tau) \simeq \mu_0^{n_1} \frac{U_{11}\left(U^{-1}\right)_{11}}{x_1} \left(e^{x_1\tau} - 1\right). \tag{S49}$$

Thus, arrive at the following final solution to the aggregation dynamics:

$$\bar{M}(\tau) \simeq 1 - \exp\left(-\mu_0\, \varepsilon \int_0^\tau \bar{P_{\rm f}}^{(1)}(\tau')d\tau'\right) \simeq 1 - \exp\left(-\frac{\mu_0^{n_1+1}\, \varepsilon\, U_{11}\left(U^{-1}\right)_{11}}{x_1^2} \left(e^{x_1\tau} - 1\right)\right). \tag{S50}$$

Transforming back to real units, we can write Eq. (S50) as

$$\boxed{\frac{M(t)}{m_{\rm tot}} = 1 - \exp\left(-\frac{\lambda_{\rm eff}^2}{2\kappa_{\rm eff}^2} \left(e^{\kappa_{\rm eff} t} - 1\right)\right)} \tag{S51}$$

where

$$\boxed{\frac{\kappa_{\rm eff}}{\kappa} = \frac{x_1}{\sqrt{\nu_2}}, \quad \frac{\lambda_{\rm eff}}{\lambda} = \sqrt{2\mu_0^{n_1+1}\, U_{11}\left(U^{-1}\right)_{11}}} \tag{S52}$$

In the following, we use Eq. (S50) to obtain explicit solutions to the aggregation kinetics in specific inhibition scenarios of practical importance.

**S2.4. Binding to fibril ends.** For the case when the inhibitor binds to the ends of fibrils, Eq. (S40) becomes

$$\frac{d}{dt}\begin{pmatrix} \bar{P_{\rm f}}^{(1)} \\ \bar{m}^{(1)} \\ \bar{P_{\rm b}}^{(1)} \end{pmatrix} = \begin{pmatrix} -\beta_{\rm e} & -\nu_2 & \alpha_{\rm e} \\ -1 & 0 & 0 \\ \beta_{\rm e} & 0 & -\alpha_{\rm e} \end{pmatrix} \begin{pmatrix} \bar{P_{\rm f}}^{(1)} \\ \bar{m}^{(1)} \\ \bar{P_{\rm b}}^{(1)} \end{pmatrix} + \begin{pmatrix} 1 \\ 0 \\ 0 \end{pmatrix}. \tag{S53}$$

The solution is given by a sum of exponential terms; to find these terms we need to consider the characteristic polynomial of the matrix in Eq. (S53) to determine its eigenvalues $x_1, x_2, x_3$:

$$x^3 + (\alpha_{\rm e} + \beta_{\rm e})x^2 - \nu_2 x - \alpha_{\rm e}\nu_2 = 0. \tag{S54}$$

There are 2 negative and 1 positive eigenvalues, which can be determined explicitly from the solution of the cubic characteristic equation

$$\frac{x_1}{\sqrt{\nu_2}} = -\frac{1}{3}\left(\gamma + C + \frac{\Delta_0}{C}\right), \tag{S55a}$$

$$\frac{x_2}{\sqrt{\nu_2}} = -\frac{1}{3}\left(\gamma + \xi C + \xi^2 \frac{\Delta_0}{C}\right), \tag{S55b}$$

$$\frac{x_3}{\sqrt{\nu_2}} = -\frac{1}{3}\left(\gamma + \xi^2 C + \xi \frac{\Delta_0}{C}\right), \tag{S55c}$$

where $\xi = \frac{-1+\sqrt{3}i}{2}$, $\gamma = \frac{\alpha_{\rm e}+\beta_{\rm e}}{\sqrt{\nu_2}}$, $C = \left(\frac{\Delta_1 + \sqrt{\Delta_1^2 - 4\Delta_0^3}}{2}\right)^{1/3}$, $\Delta_0 = \left(\frac{\gamma}{\sqrt{\nu_2}}\right)^2 + 3$, $\Delta_1 = 2\left(\frac{\gamma}{\sqrt{\nu_2}}\right)^3 - 9\frac{\gamma}{\sqrt{\nu_2}} - 27\frac{\alpha_{\rm e}}{\sqrt{\nu_2}}$. The associated eigenvectors are

$$\begin{pmatrix} \frac{\alpha_{\rm e}+x_1}{\beta_{\rm e}} \\ -\frac{\alpha_{\rm e}+x_1}{\beta_{\rm e} x_1} \\ 1 \end{pmatrix}, \quad \begin{pmatrix} \frac{\alpha_{\rm e}+x_2}{\beta_{\rm e}} \\ -\frac{\alpha_{\rm e}+x_2}{\beta_{\rm e} x_2} \\ 1 \end{pmatrix}, \quad \begin{pmatrix} \frac{\alpha_{\rm e}+x_3}{\beta_{\rm e}} \\ -\frac{\alpha_{\rm e}+x_3}{\beta_{\rm e} x_3} \\ 1 \end{pmatrix}. \tag{S56}$$

The matrix $\boldsymbol{U}$ is therefore given by

$$\boldsymbol{U} = \begin{pmatrix} \frac{\alpha_{\rm e}+x_1}{\beta_{\rm e}} & \frac{\alpha_{\rm e}+x_2}{\beta_{\rm e}} & \frac{\alpha_{\rm e}+x_3}{\beta_{\rm e}} \\ -\frac{\alpha_{\rm e}+x_1}{\beta_{\rm e} x_1} & -\frac{\alpha_{\rm e}+x_2}{\beta_{\rm e} x_2} & -\frac{\alpha_{\rm e}+x_3}{\beta_{\rm e} x_3} \\ 1 & 1 & 1 \end{pmatrix}, \tag{S57}$$

such that

$$U_{11} = \frac{\alpha_\mathrm{e} + x_1}{\beta_\mathrm{e}}, \quad \left(U^{-1}\right)_{11} = \frac{\beta_\mathrm{e} x_1}{(x_1 - x_2)(x_1 - x_3)}. \tag{S58}$$

Thus, using Eq. (S50) the solution to the aggregation kinetics is

$$\bar{M}(\tau) \simeq 1 - \exp\left(-\frac{\varepsilon U_{11}\left(U^{-1}\right)_{11}}{x_1^2}\left(e^{x_1\tau} - 1\right)\right) = 1 - \exp\left(-\frac{\varepsilon(x_1 + \alpha_\mathrm{e})}{x_1(x_1 - x_2)(x_1 - x_3)}\left(e^{x_1\tau} - 1\right)\right), \tag{S59}$$

where we used $\mu_0 = 1$ (no monomer inhibition in this case). The explicit expressions for $\kappa_\mathrm{eff}$ and $\lambda_\mathrm{eff}$ in the case that the inhibitor binds aggregate ends are obtained from Eq. (S52) as

$$\boxed{\frac{\lambda_\mathrm{eff}}{\lambda} = \sqrt{\frac{2x_1(x_1 + \alpha_\mathrm{e})}{(x_1 - x_2)(x_1 - x_3)}}} \tag{S60a}$$

$$\boxed{\frac{\kappa_\mathrm{eff}}{\kappa} = \frac{x_1}{\sqrt{\nu_2}}} \tag{S60b}$$

where $x_1$, $x_2$ and $x_3$ are given in Eq. (S55). We note that $\kappa_\mathrm{eff}/\kappa$ and $\lambda_\mathrm{eff}/\lambda$ depend only on two dimensionless combinations (Table S2):

$$a := \frac{\beta_\mathrm{e}}{\sqrt{\nu_2}} = \frac{k_\mathrm{e}^\mathrm{on} C_i}{\kappa} \quad \text{and} \quad b := \frac{\beta_\mathrm{e}}{\alpha_\mathrm{e}} = K_\mathrm{e}^\mathrm{eq} C_i. \tag{S61}$$

***S2.4.1. Dominant balance – fast binding limit.*** We have solved Eq. (S54) explicitly, but the resulting expressions (Eq. (S55)) are in general complicated. In the limit of fast binding to fibril ends, we can obtain approximate expressions for the roots of the characteristic polynomial using a dominant balance method (9) as follows. Fast binding corresponds to the situation when

$$a = \frac{\beta_\mathrm{e}}{\sqrt{\nu_2}} \gg 1 \quad \Leftrightarrow \quad \frac{k_\mathrm{e}^\mathrm{on} C_i}{\kappa} \gg 1. \tag{S62}$$

In this case, we can solve the characteristic equation Eq. (S54) considering $\sqrt{\nu_2}$ as a small parameter. The possible dominant balances for Eq. (S54) are:

- If $x = \mathcal{O}(1)$, then the dominant balance equation is $x^3 + (\alpha_\mathrm{e} + \beta_\mathrm{e})x^2 = 0$; hence $x \simeq -(\alpha_\mathrm{e} + \beta_\mathrm{e})$.
- If $x = \mathcal{O}(\sqrt{\nu_2})$, then the dominant balance equation is $(\alpha_\mathrm{e} + \beta_\mathrm{e})X^2 - \alpha = 0$, where $x = \sqrt{\nu_2}X$; hence $X \simeq \pm\sqrt{\alpha/(\alpha_\mathrm{e} + \beta_\mathrm{e})}$, i.e. $x \simeq \pm\sqrt{\alpha_\mathrm{e}\nu_2/(\alpha_\mathrm{e} + \beta_\mathrm{e})}$.

In summary, the three approximate eigenvalues found using dominant balance are:

$$x_1 = \sqrt{\frac{\alpha_\mathrm{e}\nu_2}{\alpha_\mathrm{e} + \beta_\mathrm{e}}}, \quad x_2 = -\sqrt{\frac{\alpha_\mathrm{e}\nu_2}{\alpha_\mathrm{e} + \beta_\mathrm{e}}}, \quad x_3 = -(\alpha_\mathrm{e} + \beta_\mathrm{e}). \tag{S63}$$

We have one positive and two negative eigenvalues. Using Eq. (S63) in Eq. (S74) with Eq. (S62), we find

$$\frac{U_{11}\left(U^{-1}\right)_{11}}{x_1^2} \simeq \frac{1}{2\nu_2} \quad \Rightarrow \quad \bar{M}(\tau) \simeq 1 - \exp\left(-\frac{\varepsilon}{2\nu_2}\left(e^{x_1\tau} - 1\right)\right). \tag{S64}$$

Transforming back to real units, we arrive at the final solution

$$\frac{M(t)}{m_\mathrm{tot}} = 1 - \exp\left(-\frac{\lambda_\mathrm{eff}^2}{2\kappa_\mathrm{eff}^2}\left(e^{\kappa_\mathrm{eff} t} - 1\right)\right), \tag{S65}$$

where

$$\frac{\lambda_\mathrm{eff}}{\lambda} = \left(\frac{\alpha_\mathrm{e}}{\alpha_\mathrm{e} + \beta_\mathrm{e}}\right)^{\frac{1}{2}} = \left(\frac{1}{1 + K_\mathrm{e}^\mathrm{eq} C_i^\mathrm{tot}}\right)^{\frac{1}{2}}, \tag{S66}$$

$$\frac{\kappa_\mathrm{eff}}{\kappa} = \left(\frac{\alpha_\mathrm{e}}{\alpha_\mathrm{e} + \beta_\mathrm{e}}\right)^{\frac{1}{2}} = \left(\frac{1}{1 + K_\mathrm{e}^\mathrm{eq} C_i^\mathrm{tot}}\right)^{\frac{1}{2}}. \tag{S67}$$

In summary, in the case of binding to fibril ends, the rate parameters are renormalized according to the following scheme:

$$\frac{k_+^\mathrm{eff}}{k_+} = \frac{1}{1 + K_\mathrm{e}^\mathrm{eq} C_i^\mathrm{tot}}, \tag{S68}$$

$$\frac{k_1^\mathrm{eff}}{k_1} = 1, \tag{S69}$$

$$\frac{k_2^\mathrm{eff}}{k_2} = 1. \tag{S70}$$

Our asymptotic analysis thus recovers the pre-equilibrium solution found in Sec. S1.4 in the limit of fast inhibitor binding.

**S2.5. Binding to fibril surface.** When the inhibitor binds to the surface of fibrils, Eq. (S40) becomes

$$\frac{d}{dt}\begin{pmatrix}\bar{P}^{(1)}\\ \bar{m}^{(1)}\\ \bar{M_{\rm b}}^{(1)}\end{pmatrix}=\begin{pmatrix}0 & -\nu_2 & -\nu_2\\ -1 & 0 & 0\\ 0 & -\beta_{\rm s} & -(\alpha_{\rm s}+\beta_{\rm s})\end{pmatrix}\begin{pmatrix}\bar{P}^{(1)}\\ \bar{m}^{(1)}\\ \bar{M_{\rm b}}^{(1)}\end{pmatrix}+\begin{pmatrix}1\\0\\0\end{pmatrix}. \qquad \text{[S71]}$$

To find the eigenvalues of the above matrix, we consider its characteristic polynomial:

$$x^3+(\alpha_{\rm s}+\beta_{\rm s})x^2-\nu_2 x-\alpha_{\rm s}\nu_2=0. \qquad \text{[S72]}$$

The 3 eigenvalues are given by Eq. (S55) if $\alpha_{\rm e}$ and $\beta_{\rm e}$ are replaced by $\alpha_{\rm s}$ and $\beta_{\rm s}$. The associated eigenvectors are

$$\begin{pmatrix}\frac{x_1(\alpha_{\rm s}+\beta_{\rm s}+x_1)}{\beta_{\rm s}}\\ -\frac{\alpha_{\rm s}+\beta_{\rm s}+x_1}{\beta_{\rm s}x_1}\\ 1\end{pmatrix},\quad \begin{pmatrix}\frac{x_2(\alpha_{\rm s}+\beta_{\rm s}+x_2)}{\beta_{\rm s}}\\ -\frac{\alpha_{\rm s}+\beta_{\rm s}+x_2}{\beta_{\rm s}x_2}\\ 1\end{pmatrix},\quad \begin{pmatrix}\frac{x_3(\alpha_{\rm s}+\beta_{\rm s}+x_3)}{\beta_{\rm s}}\\ -\frac{\alpha_{\rm s}+\beta_{\rm s}+x_3}{\beta_{\rm s}x_3}\\ 1\end{pmatrix}, \qquad \text{[S73]}$$

such that

$$U_{11}=\frac{x_1(\alpha_{\rm s}+\beta_{\rm s}+x_1)}{\beta_{\rm s}},\qquad \left(U^{-1}\right)_{11}=\frac{\beta_{\rm s}}{(x_1-x_2)(x_1-x_3)}. \qquad \text{[S74]}$$

Thus, using Eq. (S50) the solution to the aggregation kinetics is

$$\bar{M}(\tau)\simeq 1-\exp\left(-\frac{\varepsilon\, U_{11}\left(U^{-1}\right)_{11}}{x_1^2}\left(e^{x_1\tau}-1\right)\right)=1-\exp\left(-\frac{\varepsilon(x_1+\alpha_{\rm s}+\beta_{\rm s})}{x_1(x_1-x_2)(x_1-x_3)}\left(e^{x_1\tau}-1\right)\right). \qquad \text{[S75]}$$

The explicit expressions for $\kappa_{\rm eff}$ and $\lambda_{\rm eff}$ are obtained from Eq. (S52) as

$$\boxed{\frac{\lambda_{\rm eff}}{\lambda}=\sqrt{\frac{2x_1(x_1+\alpha_{\rm s}+\beta_{\rm s})}{(x_1-x_2)(x_1-x_3)}}} \qquad \text{[S76a]}$$

$$\boxed{\frac{\kappa_{\rm eff}}{\kappa}=\frac{x_1}{\sqrt{\nu_2}}} \qquad \text{[S76b]}$$

where $x_1$, $x_2$ and $x_3$ are given in Eq. (S55) by replacing $\alpha_{\rm e}$ and $\beta_{\rm e}$ by $\alpha_{\rm s}$ and $\beta_{\rm s}$. Thus, in this case the relevant dimensionless combinations of parameters that determine $\kappa_{\rm eff}$ and $\lambda_{\rm eff}$ are (Table S2):

$$a:=\frac{\beta_{\rm s}}{\sqrt{\nu_2}}=\frac{k_{\rm s}^{\rm on}C_i}{\kappa}\quad\text{and}\quad b:=\frac{\beta_{\rm s}}{\alpha_{\rm s}}=K_{\rm s}^{\rm eq}C_i. \qquad \text{[S77]}$$

***S2.5.1. Dominant balance – fast inhibitor binding limit.*** In the limit of fast binding to fibril surfaces, $\sqrt{\nu_2}\ll\alpha_{\rm s},\beta_{\rm s}$, we can obtain approximate expressions for the roots of the characteristic polynomial using a dominant balance argument as above. The eigenvalues are approximatively given by

$$x_1=\sqrt{\frac{\alpha_{\rm s}\nu_2}{\alpha_{\rm s}+\beta_{\rm s}}},\quad x_2=-\sqrt{\frac{\alpha_{\rm s}\nu_2}{\alpha_{\rm s}+\beta_{\rm s}}},\quad x_3=-(\alpha_{\rm s}+\beta_{\rm s}) \qquad \text{[S78]}$$

Thus,

$$\frac{U_{11}\left(U^{-1}\right)_{11}}{x_1^2}\simeq\frac{1}{2x_1^2}\quad\Rightarrow\quad \bar{M}(\tau)\simeq 1-\exp\left(-\frac{\varepsilon}{2x_1^2}\left(e^{x_1\tau}-1\right)\right). \qquad \text{[S79]}$$

Transforming back to real units, we arrive at the final solution

$$\frac{M(t)}{m_{\rm tot}}=1-\exp\left(-\frac{\lambda_{\rm eff}^2}{2\kappa_{\rm eff}^2}\left(e^{\kappa_{\rm eff}t}-1\right)\right), \qquad \text{[S80]}$$

where

$$\frac{\lambda_{\rm eff}}{\lambda}=1, \qquad \text{[S81]}$$

$$\frac{\kappa_{\rm eff}}{\kappa}=\left(\frac{\alpha_{\rm s}}{\alpha_{\rm s}+\beta_{\rm s}}\right)^{\frac{1}{2}}=\left(\frac{1}{1+K_{\rm s}^{\rm eq}C_i^{\rm tot}}\right)^{\frac{1}{2}}. \qquad \text{[S82]}$$

In summary, in the case of fast inhibitor binding to the fibril surface, the renormalized rate parameters are given by:

$$\frac{k_+^{\rm eff}}{k_+}=1, \qquad \text{[S83]}$$

$$\frac{k_1^{\rm eff}}{k_1}=1, \qquad \text{[S84]}$$

$$\frac{k_2^{\rm eff}}{k_2}=\frac{1}{1+K_{\rm s}^{\rm eq}C_i^{\rm tot}}, \qquad \text{[S85]}$$

which is the same result of Sec. S1.4.

**S2.6. Binding to monomers.** When the inhibitor binds to monomers, Eq. (S40) becomes

$$\frac{d}{dt}\begin{pmatrix} \bar{P_\mathrm{f}}^{(1)} \\ \bar{m_\mathrm{f}}^{(1)} \\ \bar{m_\mathrm{b}}^{(1)} \end{pmatrix} = \begin{pmatrix} 0 & -\nu_2\mu_0^{n_2} & -\nu_2\mu_0^{n_2} \\ -\mu_0 & -\beta_\mathrm{m} & \alpha_\mathrm{m} \\ 0 & \beta_\mathrm{m} & -\alpha_\mathrm{m} \end{pmatrix}\begin{pmatrix} \bar{P_\mathrm{f}}^{(1)} \\ \bar{m_\mathrm{f}}^{(1)} \\ \bar{m_\mathrm{b}}^{(1)} \end{pmatrix} + \begin{pmatrix} \mu_0^{n_1} \\ 0 \\ 0 \end{pmatrix}, \tag{S86}$$

The eigenvalues of the above matrix are

$$x_1 = \sqrt{\nu_2\mu_0^{n_2+1}}, \quad x_2 = -\sqrt{\nu_2\mu_0^{n_2+1}}, \quad x_3 = -(\alpha_\mathrm{m}+\beta_\mathrm{m}) \tag{S87}$$

with associated eigenvectors

$$\begin{pmatrix} -\frac{x_1(\alpha_\mathrm{m}+\beta_\mathrm{m}+x_1)}{\beta_\mathrm{m}\mu_0} \\ \frac{\alpha_\mathrm{m}+x_1}{\beta_\mathrm{m}} \\ 1 \end{pmatrix}, \quad \begin{pmatrix} -\frac{x_2(\alpha_\mathrm{m}+\beta_\mathrm{m}+x_2)}{\beta_\mathrm{m}\mu_0} \\ \frac{\alpha_\mathrm{m}+x_2}{\beta_\mathrm{m}} \\ 1 \end{pmatrix}, \quad \begin{pmatrix} 0 \\ -1 \\ 1 \end{pmatrix} \tag{S88}$$

It follows

$$U_{11} = -\frac{x_1(\alpha_\mathrm{m}+\beta_\mathrm{m}+x_1)}{\beta_\mathrm{m}\mu_0} \qquad \left(U^{-1}\right)_{11} = -\frac{\beta_\mathrm{m}\mu_0}{2x_1(\alpha_\mathrm{m}+\beta_\mathrm{m}+x_1)} \tag{S89}$$

Using Eq. (S52), we find the following expressions for the effective $\lambda$ and $\kappa$

$$\boxed{\frac{\lambda_\mathrm{eff}}{\lambda} = \sqrt{2\mu_0^{n_1+1}\,U_{11}\left(U^{-1}\right)_{11}} = \mu_0^{\frac{n_1+1}{2}}} \tag{S90a}$$

$$\boxed{\frac{\kappa_\mathrm{eff}}{\kappa} = \frac{x_1}{\sqrt{\nu_2}} = \mu_0^{\frac{n_2+1}{2}}} \tag{S90b}$$

**S2.7. Combining binding to monomers, fibril ends and fibril surface.** We now consider the case when the inhibitor can bind all protein species, i.e. monomers, fibril ends and surfaces. We need to solve

$$\frac{d}{dt}\begin{pmatrix} \bar{P_\mathrm{f}}^{(1)} \\ \bar{m_\mathrm{f}}^{(1)} \\ \bar{m_\mathrm{b}}^{(1)} \\ \bar{P_\mathrm{b}}^{(1)} \\ \bar{M_\mathrm{b}}^{(1)} \end{pmatrix} = \begin{pmatrix} -\beta_\mathrm{e} & -\nu_2\mu_0^{n_2} & -\nu_2\mu_0^{n_2} & \alpha_\mathrm{e} & -\nu_2\mu_0^{n_2} \\ -\mu_0 & -\beta_\mathrm{m} & \alpha_\mathrm{m} & 0 & 0 \\ 0 & \beta_\mathrm{m} & -\alpha_\mathrm{m} & 0 & 0 \\ \beta_\mathrm{e} & 0 & 0 & -\alpha_\mathrm{e} & 0 \\ 0 & -\beta_\mathrm{s} & -\beta_\mathrm{s} & 0 & -(\alpha_\mathrm{s}+\beta_\mathrm{s}) \end{pmatrix}\begin{pmatrix} \bar{P_\mathrm{f}}^{(1)} \\ \bar{m_\mathrm{f}}^{(1)} \\ \bar{m_\mathrm{b}}^{(1)} \\ \bar{P_\mathrm{b}}^{(1)} \\ \bar{M_\mathrm{b}}^{(1)} \end{pmatrix} + \begin{pmatrix} \mu_0^{n_1} \\ 0 \\ 0 \\ 0 \\ 0 \end{pmatrix}, \tag{S91}$$

where $\mu_0 = \alpha_m/(\alpha_m+\beta_m)$. The characteristic polynomial of the above matrix is

$$(x+\alpha_m+\beta_m)\Big[x^4 + (\alpha_e+\beta_e+\alpha_s+\beta_s)x^3 + [(\alpha_e+\beta_e)(\alpha_s+\beta_s) - \nu_2\mu_0^{n_2+1}]x^2 - (\alpha_e+\alpha_s)\nu_2\mu_0^{n_2+1}x - \alpha_e\alpha_s\nu_2\mu_0^{n_2+1}\Big] = 0. \tag{S92}$$

Using dominant balance argument, we can find an approximated expression for the largest (positive) eigenvalue $x_1$ by writing $x_1 = \sqrt{\nu_2}X_1$ leading to $(\alpha_e+\beta_e)(\alpha_s+\beta_s)X_1^2 - \alpha_e\alpha_s\mu_0^{n_2+1} = 0$, i.e.

$$x_1 \simeq \sqrt{\nu_2\left(\frac{\alpha_e}{\alpha_e+\beta_e}\right)\left(\frac{\alpha_s}{\alpha_s+\beta_s}\right)\left(\frac{\alpha_m}{\alpha_m+\beta_m}\right)^{n_2+1}}. \tag{S93}$$

Using Eq. (S50) the final solution for the aggregate mass is found to be:

$$\frac{M(t)}{m(0)} = 1 - \exp\left(-\frac{\lambda_\mathrm{eff}^2}{2\kappa_\mathrm{eff}^2}\left[e^{\kappa_\mathrm{eff}t} - 1\right]\right), \tag{S94}$$

where

$$\frac{\lambda_\mathrm{eff}}{\lambda} = \left(\frac{\alpha_\mathrm{e}}{\alpha_\mathrm{e}+\beta_\mathrm{e}}\right)^{\frac{1}{2}}\left(\frac{\alpha_\mathrm{m}}{\alpha_\mathrm{m}+\beta_\mathrm{m}}\right)^{\frac{n_1+1}{2}} = \left(\frac{1}{1+K_\mathrm{m}^\mathrm{eq}C_i^\mathrm{tot}}\right)^{\frac{n_1+1}{2}}\left(\frac{1}{1+K_\mathrm{e}^\mathrm{eq}C_i^\mathrm{tot}}\right)^{\frac{1}{2}}, \tag{S95}$$

$$\frac{\kappa_\mathrm{eff}}{\kappa} = \left(\frac{\alpha_\mathrm{e}}{\alpha_\mathrm{e}+\beta_\mathrm{e}}\right)^{\frac{1}{2}}\left(\frac{\alpha_\mathrm{s}}{\alpha_\mathrm{s}+\beta_\mathrm{s}}\right)^{\frac{1}{2}}\left(\frac{\alpha_\mathrm{m}}{\alpha_\mathrm{m}+\beta_\mathrm{m}}\right)^{\frac{n_2+1}{2}} = \left(\frac{1}{1+K_\mathrm{m}^\mathrm{eq}C_i^\mathrm{tot}}\right)^{\frac{n_2+1}{2}}\left(\frac{1}{1+K_\mathrm{e}^\mathrm{eq}C_i^\mathrm{tot}}\right)^{\frac{1}{2}}\left(\frac{1}{1+K_\mathrm{s}^\mathrm{eq}C_i^\mathrm{tot}}\right)^{\frac{1}{2}}. \tag{S96}$$

**Table S2.** Effective rates of aggregation $\frac{\lambda^{\text{eff}}}{\lambda}$ and $\frac{\kappa^{\text{eff}}}{\kappa}$ as a function of $a = k_{\times}^{\text{on}} C_i/\kappa$ and $b = K_{\times}^{\text{eq}} C_i$. These functions yield the plots in Fig. 3a,b of the main text. In this table $y_1$, $y_2$, $y_3$ are the 3 roots of the equation $y^3 + \left(a + \frac{a}{b}\right) y^2 - y - \frac{a}{b} = 0$.

| Targeted species | Monomers | Aggregate ends | Fibril surface |
|---|---|---|---|
| Effective rate of 1. pathways $\frac{\lambda^{\text{eff}}}{\lambda}$ | $\left(\frac{1-e^{-\frac{\kappa}{\lambda}\left(a+\frac{a}{b}\right)}}{1+b}\right)^{\frac{n_1+1}{2}}$ | $\left(\frac{2y_1\left(y_1+\frac{a}{b}\right)}{(y_1-y_2)(y_1-y_3)}\right)^{\frac{1}{2}}$ | 1 |
| Effective rate of 2. pathways $\frac{\kappa^{\text{eff}}}{\kappa}$ | $\left(\frac{1-e^{-\left(a+\frac{a}{b}\right)}}{1+b}\right)^{\frac{n_2+1}{2}}$ | $y_1$ | $y_1$ |
| Parameter definition | $y_1 = -\frac{1}{3}\left(a + \frac{a}{b} + C + \frac{\Delta_0}{C}\right)$ | $y_2 = -\frac{1}{3}\left(a + \frac{a}{b} + \xi C + \xi^2 \frac{\Delta_0}{C}\right)$ | $y_3 = -\frac{1}{3}\left(a + \frac{a}{b} + \xi^2 C + \xi \frac{\Delta_0}{C}\right)$ |
| | $a = \frac{k_{\times}^{\text{on}} C_i}{\kappa}$, $b = K_{\times}^{\text{eq}} C_i$ | $\gamma = a + \frac{a}{b}$ | $\xi = \frac{-1+\sqrt{3}i}{2}$ |
| | $C = \left(\frac{\Delta_1+\sqrt{\Delta_1^2-4\Delta_0^3}}{2}\right)^{1/3}$ | $\Delta_0 = \left(a + \frac{a}{b}\right)^2 + 3$ | $\Delta_1 = 2\left(a + \frac{a}{b}\right)^3 - 9\left(a + \frac{a}{b}\right) - 27\frac{a}{b}$ |

In summary, our asymptotic analysis of Eq. (S12) shows that, in the case when the inhibitor can bind monomers, fibril ends and surfaces, the rate parameters are renormalized according to the following scheme:

$$\frac{k_+^{\text{eff}}}{k_+} = \left(\frac{1}{1 + K_{\text{m}}^{\text{eq}} C_i^{\text{tot}}}\right)\left(\frac{1}{1 + K_{\text{e}}^{\text{eq}} C_i^{\text{tot}}}\right), \tag{S97}$$

$$\frac{k_1^{\text{eff}}}{k_1} = \left(\frac{1}{1 + K_{\text{m}}^{\text{eq}} C_i^{\text{tot}}}\right)^{n_1}, \tag{S98}$$

$$\frac{k_2^{\text{eff}}}{k_2} = \left(\frac{1}{1 + K_{\text{m}}^{\text{eq}} C_i^{\text{tot}}}\right)^{n_2}\left(\frac{1}{1 + K_{\text{s}}^{\text{eq}} C_i^{\text{tot}}}\right), \tag{S99}$$

which recovers Eq. (S23). The effect of the individual modes of inhibition on $\lambda$ and $\kappa$ combine multiplicatively.

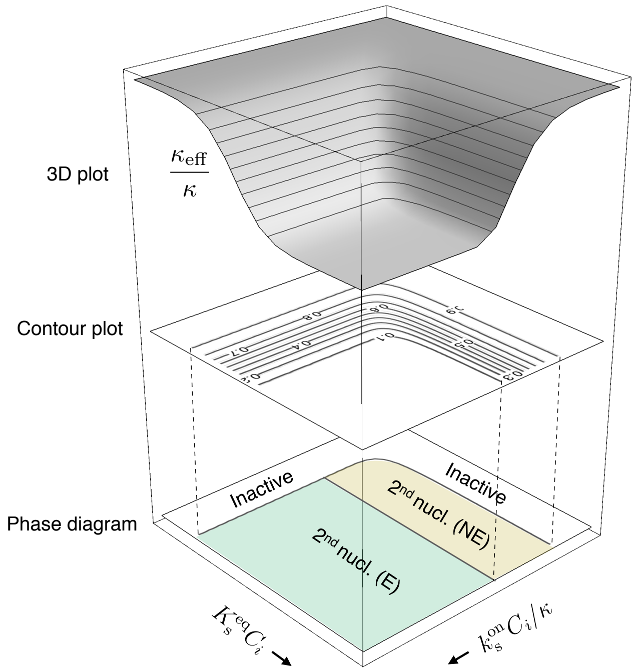

**Fig. S3.** Construction of phase diagram of possible inhibition regimes for an inhibitor that binds fibril surface sites. Contour lines are shown here in steps of 0.1. Boundary lines are however not sharp: the extent of inhibition is in fact a continuous function of $a = k_{\times}^{\text{on}} C_i/\kappa$ and $b = K_{\times}^{\text{eq}} C_i$.

### S3. Asymptotic solutions to aggregation kinetics with variable inhibitor concentration

Our asymptotic approach for solving Eq. (S12) can be generalized straightforwardly to account for the situation of variable binder concentration. We discuss here this idea on the example of monomer binding; the treatment for binding to fibril ends or fibril surfaces is fully analogous. On accounting for a variable binder concentration, the nondimensionalized moment equations in this case are:

$$\frac{\mathrm{d}\bar{P}(\tau)}{\mathrm{d}\tau} = \varepsilon\, \bar{m}_\mathrm{f}(\tau)^{n_1} + \nu_2\, \bar{m}_\mathrm{f}(\tau)^{n_2}\Big(1 - \bar{m}_\mathrm{f}(\tau) - \bar{m}_\mathrm{b}(\tau)\Big), \tag{S100a}$$

$$\frac{\mathrm{d}\bar{m}_\mathrm{f}(\tau)}{\mathrm{d}\tau} = -\bar{m}_\mathrm{f}(\tau)\bar{P}(\tau) - \beta\, \bar{C}_i(\tau)\bar{m}_\mathrm{f}(\tau) + \alpha\, \bar{m}_\mathrm{b}(\tau), \tag{S100b}$$

$$\frac{\mathrm{d}\bar{m}_\mathrm{b}(\tau)}{\mathrm{d}\tau} = \beta\, \bar{C}_i(\tau)\bar{m}_\mathrm{f}(\tau) - \alpha\, \bar{m}_\mathrm{b}(\tau) = -\frac{\mathrm{d}\bar{C}_i(\tau)}{\mathrm{d}\tau}, \tag{S100c}$$

where

$$\bar{C}_i = \frac{C_i}{m_\mathrm{tot}}, \tag{S100d}$$

$$\beta = \frac{k_\mathrm{m}^\mathrm{on}}{2k_+ m_\mathrm{tot}}, \tag{S100e}$$

$$\alpha = \frac{k_\mathrm{m}^\mathrm{off}}{2k_+ m_\mathrm{tot}}. \tag{S100f}$$

Using a naive perturbation expansion $\bar{P} = \bar{P}^{(0)} + \varepsilon \bar{P}^{(0)} + \cdots$, etc., the equations at order $\varepsilon^0$ are found to be:

$$\frac{\mathrm{d}\bar{P}^{(0)}(\tau)}{\mathrm{d}\tau} = \nu_2\, \bar{m}_\mathrm{f}{}^{(0)}(\tau)^{n_2}\Big(1 - \bar{m}_\mathrm{f}{}^{(0)}(\tau) - \bar{m}_\mathrm{b}{}^{(0)}(\tau)\Big), \tag{S101a}$$

$$\frac{\mathrm{d}\bar{m}_\mathrm{f}{}^{(0)}(\tau)}{\mathrm{d}\tau} = -\bar{m}_\mathrm{f}{}^{(0)}(\tau)\bar{P}^{(0)}(\tau) - \beta\, \bar{C}_i{}^{(0)}(\tau)\bar{m}_\mathrm{f}{}^{(0)}(\tau) + \alpha\, \bar{m}_\mathrm{b}{}^{(0)}(\tau), \tag{S101b}$$

$$\frac{\mathrm{d}\bar{m}_\mathrm{b}{}^{(0)}(\tau)}{\mathrm{d}\tau} = \beta\, \bar{C}_i{}^{(0)}(\tau)\bar{m}_\mathrm{f}{}^{(0)}(\tau) - \alpha\, \bar{m}_\mathrm{b}{}^{(0)}(\tau) = -\frac{\mathrm{d}\bar{C}_i{}^{(0)}(\tau)}{\mathrm{d}\tau}. \tag{S101c}$$

The solution, subject to initial conditions $\bar{m}_\mathrm{f}{}^{(0)}(0) = 1$, $\bar{C}_i{}^{(0)} = \gamma_0$, $\bar{P}^{(0)}(0) = \bar{m}_\mathrm{f}{}^{(0)}(0) = 0$, is:

$$\bar{P}^{(0)} \equiv 0, \tag{S102}$$

$$\bar{m}_\mathrm{b}{}^{(0)} = \frac{A_1 A_2\left(1 - e^{\beta\xi\tau}\right)}{A_2 - A_1 e^{\beta\xi\tau}}, \tag{S103}$$

$$\bar{m}_\mathrm{f}{}^{(0)} = 1 - \frac{A_1 A_2\left(1 - e^{\beta\xi\tau}\right)}{A_2 - A_1 e^{\beta\xi\tau}}, \tag{S104}$$

$$\bar{C}_i{}^{(0)} = \gamma_0 - \bar{m}_\mathrm{b}{}^{(0)}, \tag{S105}$$

where

$$A_{1,2} = \frac{1 + \gamma_0 + \frac{\alpha}{\beta}}{2} \pm \sqrt{\left(\frac{1 + \gamma_0 + \frac{\alpha}{\beta}}{2}\right)^2 - \gamma_0} \tag{S106}$$

and $\xi = A_1 - A_2$. Thus, we have separation of timescales and the solution is given by

$$\frac{M(t)}{m_\mathrm{tot}} = 1 - \exp\left(-\frac{\lambda_\mathrm{eff}^2}{2\kappa_\mathrm{eff}^2}\left(e^{\kappa_\mathrm{eff} t} - 1\right)\right), \tag{S107}$$

where

$$\frac{\lambda_\mathrm{eff}}{\lambda} = (1 - A_2)^{\frac{n_1+1}{2}} = \left(\frac{1}{1 + K_\mathrm{m}^\mathrm{eq} C_i^\mathrm{eq}}\right)^{\frac{n_1+1}{2}}, \tag{S108}$$

$$\frac{\kappa_\mathrm{eff}}{\kappa} = (1 - A_2)^{\frac{n_2+1}{2}} = \left(\frac{1}{1 + K_\mathrm{m}^\mathrm{eq} C_i^\mathrm{eq}}\right)^{\frac{n_2+1}{2}}. \tag{S109}$$

Hence, $1 - A_2$ corresponds to the equilibrium concentration of free monomers. The kinetics are expressed in terms effective rate parameters which are renormalized by the presence of inhibitor according to the following scheme:

$$\frac{k_+^\mathrm{inhibition}}{k_+} = \frac{1}{1 + K_\mathrm{m}^\mathrm{eq} C_i^\mathrm{eq}}, \tag{S110}$$

$$\frac{k_n^\mathrm{inhibition}}{k_n} = \left(\frac{1}{1 + K_\mathrm{m}^\mathrm{eq} C_i^\mathrm{eq}}\right)^{n_1}, \tag{S111}$$

$$\frac{k_2^\mathrm{inhibition}}{k_2} = \left(\frac{1}{1 + K_\mathrm{m}^\mathrm{eq} C_i^\mathrm{eq}}\right)^{n_2}, \tag{S112}$$

where $C_i^\mathrm{eq}$ is the equilibrium concentration of inhibitor.
